# Topoisomerase IIIα resolves inter- and intra-molecular intertwines during DNA replication

**DOI:** 10.64898/2026.06.30.735647

**Authors:** Sabrina X Van Ravenstein, Lillian V Campos, Gabor M Harami, Darren R Heintzman, Steven N Dahmen, William G Dunphy, Keir C Neuman, James M. Dewar

## Abstract

Resolution of topological stress is crucial for genome integrity. Vertebrate topoisomerase IIα (TOP2α) resolves catenanes to relieve topological stress and unlink daughter molecules during DNA replication. Topoisomerase IIIα (TOP3α) can also resolve catenanes, but its direct role during DNA replication, substrate specificity, and relevant binding partners remain unclear. Here we show that TOP3α becomes crucial when TOP2α function is compromised. We find that in *Xenopus* egg extracts, TOP3α promotes replication fork progression and daughter strand unlinking specifically during replication termination. TOP3α can carry out this role independently of its binding partners RMI1-RMI2 and the BLM helicase by acting on lagging-strand single-stranded DNA (ssDNA). Strikingly, elevated lagging-strand ssDNA drives formation of intramolecular ssDNA intertwines, which are ordinarily resolved by TOP3α. Thus, TOP3α resolves intermolecular linkages to promote fork progression during termination and resolves intramolecular linkages that arise when high levels of ssDNA are present during DNA replication.

## Introduction

DNA replication generates topological stress that manifests as supercoils ahead of the fork and pre-catenanes behind the fork (*1–4*). These structures must be quickly resolved by topoisomerases to avoid buildup of topological stress that would otherwise stall DNA replication (*5–15*). Pre-catenanes that persist after replication is complete (‘catenanes’) (*16, 17*) link the daughter DNA molecules and must be removed for faithful chromosome segregation (*18*). During eukaryotic DNA replication topoisomerase I (TOP1) can resolve supercoils (*19, 20*), while topoisomerase II (TOP2) can resolve supercoils and pre-catenanes (*21*). In contrast, the role of topoisomerase III is much less clear (*22–24*). Addressing its role is important because topoisomerase function is essential for cell viability and targeted during cancer therapy, and because TOP3A is itself a human disease gene (*25, 26*).

Topoisomerase III acts by creating a transient single-stranded DNA break through which an adjacent DNA strand, or DNA duplex, is passed (*27–30*). This activity is crucial to resolve hemicatenanes that arise from homologous recombination (*22, 31*). Bacterial topoisomerase III (topo III) can resolve catenanes (*32*) and can operate during elongation (*9, 33*) and termination (*27, 34, 35*). However, yeast topoisomerase III (Top3) cannot act during termination (*36*) and has no defined role in unperturbed replication other than at sites of fork stalling (*37*). The vertebrate replicative topoisomerase III, TOP3α, is important for DNA replication (*38*) but also resolves transcription-replication collisions (*39*) and is required for mitochondrial DNA segregation (*40–43*). It therefore remains to be established whether TOP3α promotes fork progression directly or indirectly by resolving transcription-replication collisions or ensuring mitochondrial DNA segregation.

The mechanism of topoisomerase III function during replication remains uncertain (Fig. S1A). Topoisomerase III may resolve pre-catenanes by acting on single-stranded DNA within the lagging strand template (*33*), but this has not been directly tested and is unexplored for the eukaryotic topoisomerase III paralogs. Single-stranded DNA can also arise on the leading strand template when the leading strand polymerase stalls but the replicative helicase continues to unwind (‘uncoupling’) (*44, 45*), which is frequent (*46, 47*) and thus could also be the substrate topoisomerase III acts upon (as noted in (*24*)). Consistent with this, TOP3α accumulates on DNA when uncoupling is induced (*48*). Topoisomerase III may also resolve pre-catenanes by acting on double-stranded DNA, which it can also act on (*49*), although less efficiently than single-stranded DNA (*50*). Beyond questions about substrate specificity, eukaryotic Top3/ TOP3α also interacts with additional proteins to form distinct complexes that direct its nuclear (*27, 51, 52*), but not mitochondrial (*40*), function and whose roles are incompletely understood, and TOP3α can also function non-enzymatically within these complexes (*53, 54*). For TOP3α these are the non-catalytic RMI1-RMI2 to form the ‘TRR’ complex (*55, 56*) and the BLM helicase to form the ‘BTRR’ complex (*57*). Finally, topoisomerase III can resolve intramolecular DNA linkages (*50, 58, 59*) but it is unclear whether this activity operates during replication. Thus, the substrate for topoisomerase III during DNA replication, the complexes that direct it, and the range of linkages it can resolve are all open questions.

Here, we characterized the role of TOP3α during vertebrate DNA replication using *Xenopus* egg extracts (*60, 61*). This approach allowed us to perform a detailed biochemical analysis of TOP3α using a complete vertebrate nuclear proteome but independent of transcription and mitochondrial DNA replication. We show that TOP3α normally plays no detectable role during DNA replication but promotes fork merger and pre-catenane resolution when TOP2α activity is lost. This function is independent of both RMI1-RMI2 and BLM, in contrast to previously reported nuclear functions of TOP3α. Mechanistically, TOP3α operates primarily during termination, which has not previously been reported for other eukaryotic topoisomerases. We also provide direct evidence that TOP3α acts on lagging strand gaps to resolve pre-catenanes. Additionally, we find that high levels of lagging-strand single-stranded DNA drive formation of intramolecular single-stranded DNA intertwines, which TOP3α resolves. Overall, our data reveal that TOP3α resolves intermolecular DNA intertwines (pre-catenanes) to compensate for loss of TOP2α function while playing a unique role in resolving intramolecular intertwines.

## Results

### No role for TOP3α during unperturbed DNA synthesis

To investigate the role of topoisomerase IIIα (TOP3α) during vertebrate DNA replication, we first assessed whether loss of TOP3α impacted DNA replication in *Xenopus* egg extracts (‘extracts’). Plasmid DNA was replicated in extracts depleted of TOP3α and radiolabeled nucleotides were included to monitor the extent of DNA synthesis and the products formed (Fig. S1D). Immunodepletion was effective and removed ∼>99% of TOP3α (Fig. S1E). In mock immunodepleted extracts, replication fork structures were initially detected (θs; Fig. S1F, lane 1) then converted to nicked and supercoiled circular monomeric products of DNA replication (nCMs, scCMs; Fig. S1F, lane 2) coincident with completion of DNA synthesis (Fig. S1G). These replication products were supercoiled because the plasmid DNA template is chromatinized in extracts (*62*) and the chromatin is replicated (*63*), so removal of nucleosomes results in compensatory supercoiling when the reaction is stopped (*64*). Importantly, TOP3α immunodepletion did not affect the DNA species formed (Fig. S1F), the rate of DNA synthesis (Fig. S1G), or the time taken to form circular monomeric products of DNA replication (Fig. S1H). Thus, TOP3α does not ordinarily play a role during DNA replication in *Xenopus* egg extracts.

### A TOP3α complex facilitates fork merger in the absence of TOP2α

Topoisomerase IIα () is important for catenane resolution during vertebrate DNA replication, but a low level of unlinking still takes place even in the absence of TOP2α (Fig. S1B) (*6*). We therefore investigated whether TOP3α, whose orthologs can also resolve catenanes (*49, 50*), might play a role in the absence of TOP2α (Fig. S1C). To do so, we included a catalytic inhibitor of TOP2 activity (TOP2-i, ICRF-193; characterized previously in this system (*6*)) and once again replicated DNA in either mock- or TOP3α-immunodepleted extracts (Fig. 1A). In mock-treated extracts, replication fork structures accumulated and persisted for tens of minutes (θs; Fig. 1A-B) before being converted to catenanes (cats; Fig. 1B) and a small population of high molecular weight species (HMws; Fig. 1B) as previously described. TOP3α-immunodepletion did not affect incorporation of radiolabeled nucleotides, indicating that the early stages of replication (initiation and elongation) were unaffected (Fig. 1C). However, replication fork structures persisted to the end of the reaction (Fig. 1B, lane 8). This persistence indicated that fork merger, which corresponds to complete unwinding of the parental DNA (Fig. 1A), was severely inhibited (Fig. 1D). Importantly, addition of recombinant TOP3α as part of the TOP3α-RMI1-RMI2 complex (TRR; Fig. S1I) rescued the fork merger defect (Fig. 1B, Fig. 1D), demonstrating that TOP3α or some other part of the TRR complex (see Fig. 2) promotes fork merger. Overall, these data indicate that TOP3α is required for residual fork merger upon TOP2α inhibition.

**Figure 1:**
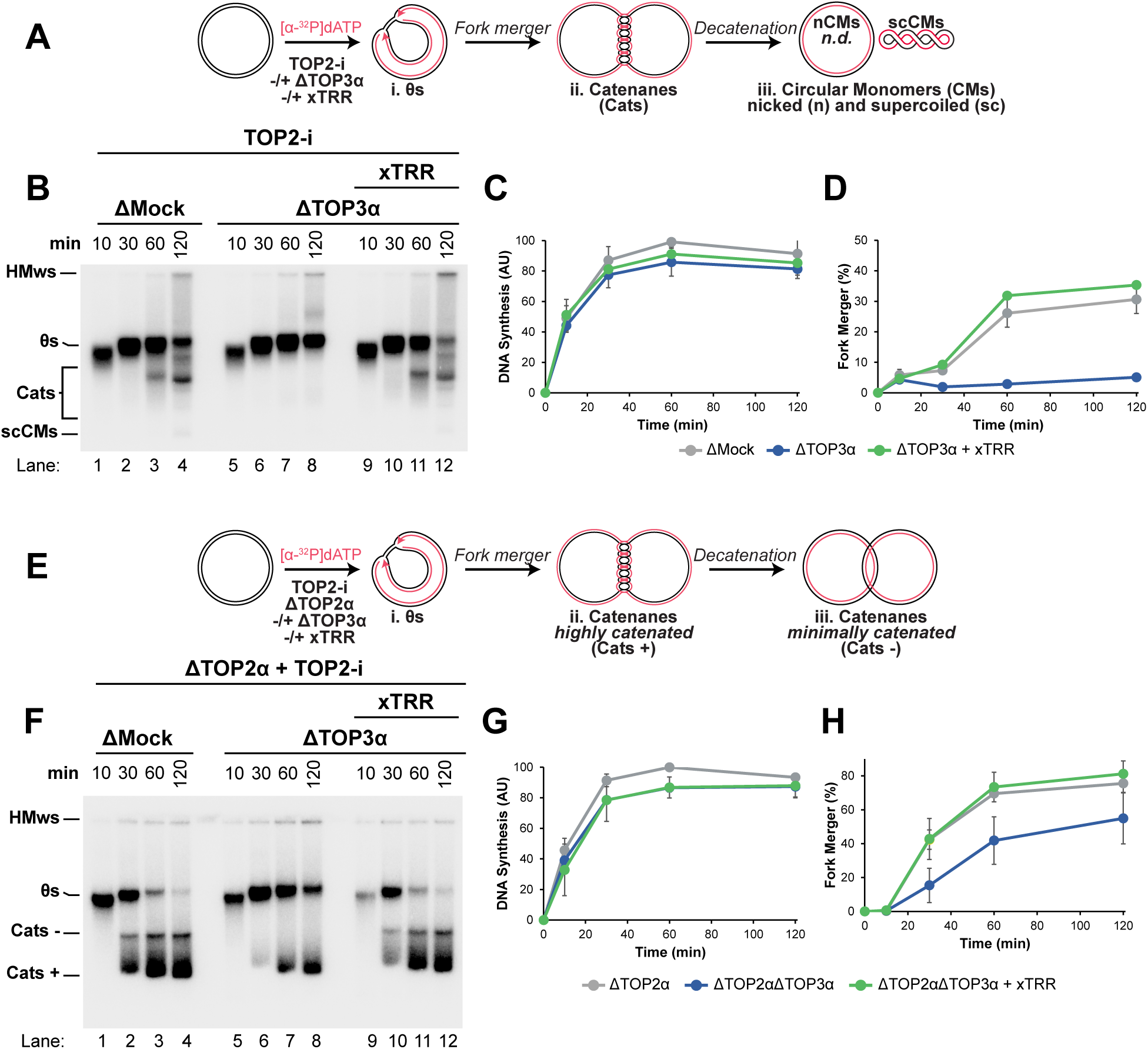
TOP3α promotes fork merger in the absence of TOP2α. (A) Plasmid DNA was replicated using *Xenopus* egg extracts in the presence of [α-^32^P]dATP to label newly-synthesized DNA strands under TOP2-inhibited (TOP2-i, ICRF-193 treated) conditions, with or without TOP3α immunodepletion and addition of recombinant *Xenopus* Topoisomerase IIIα-RMI1-RMI2 (xTRR) complex. The expected DNA structures (identified in (*6, 81*)) are indicated. Note that nicked circular monomers (nCMs) are not detected (*nd*) in this assay due to low abundance and co-migration with catenated species. (B) Replication was performed as in (A). Samples were separated on a native agarose gel and visualized by autoradiography. See also Supplemental Fig. S1D,I. (C) Quantification of total DNA synthesis from (B). (D) Quantification of fork merger from (B). (E) Plasmid DNA was replicated using *Xenopus* egg extracts in the presence of [α-^32^P]dATP to label newly-synthesized DNA strands under TOP2-immunodepleted and inhibited (TOP2-i) conditions, with or without TOP3α immunodepletion and addition of recombinant *Xenopus* Topoisomerase IIIα-RMI1-RMI2 (xTRR) complex. (F) Replication was performed as in (E). Samples were separated on a native agarose gel and visualized by autoradiography. (G) Quantification of total DNA synthesis from (F). (H) Quantification of fork merger from (F).

**Figure 2:**
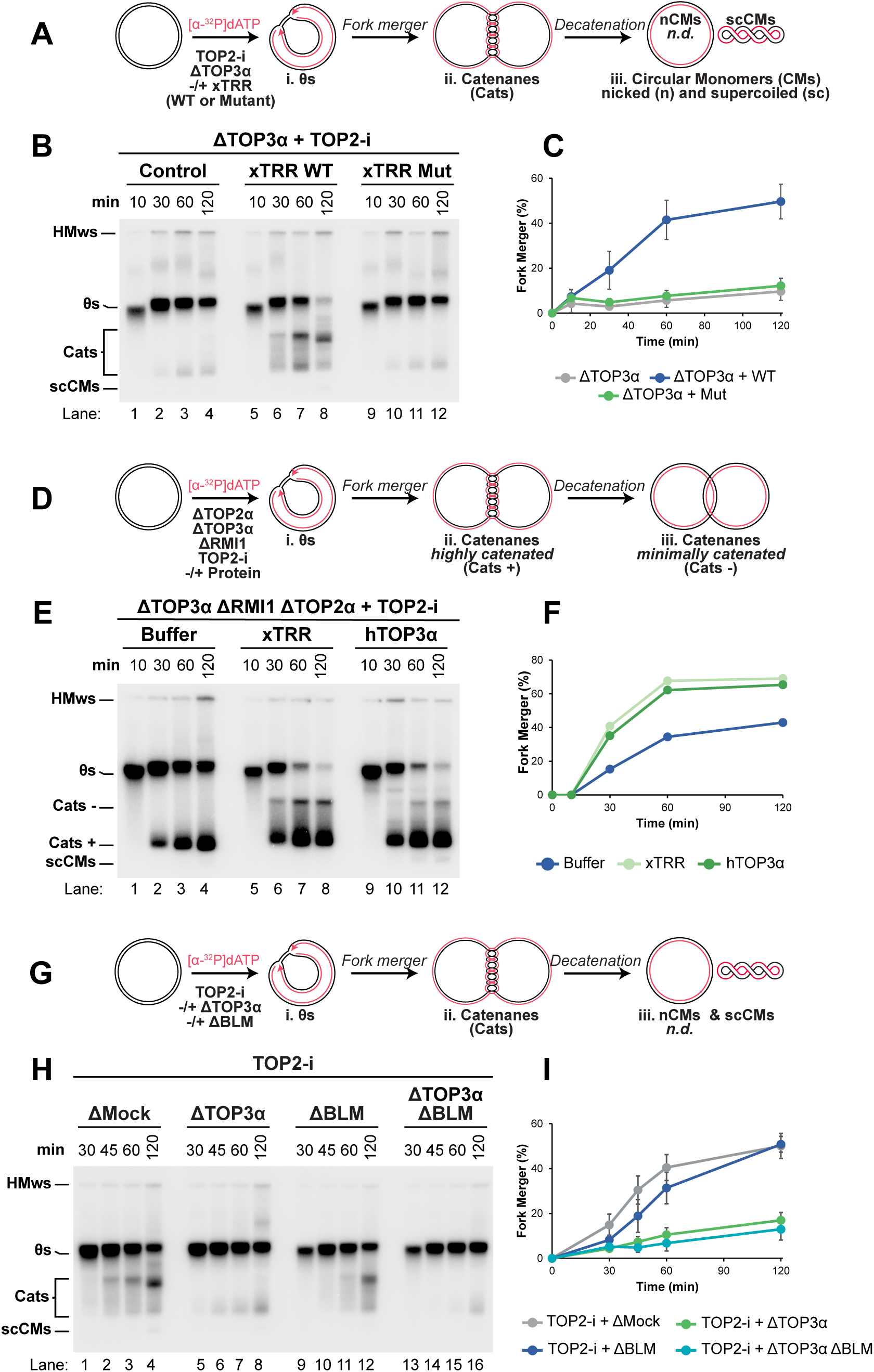
TOP3α promotes fork merger independently of RMI1-RMI2 and BLM. (A) Plasmid DNA was replicated using *Xenopus* egg extracts in the presence of [α-^32^P]dATP to label newly-synthesized DNA strands under TOP2-inhibited (TOP2-i) and TOP3α immunodepleted conditions supplemented with wild type (WT) xTRR complex or catalytic mutant (Mut). (B) Replication was performed as in (A). Samples were separated on a native agarose gel and visualized by autoradiography. See also Supplemental Fig. S2A-B. (C) Quantification of fork merger from (B). (D) Plasmid DNA was replicated using *Xenopus* egg extracts in the presence of [α-^32^P]dATP to label newly-synthesized DNA strands under TOP2 inhibited (TOP2-i) conditions co-depleted of TOP2α, TOP3α, and RMI1 with addition of recombinant xTRR complex or human TOP3α (hTOP3α). (E) Replication was performed as in (D). Samples were separated on a native agarose gel and visualized by autoradiography. (F) Quantification of fork merger from (E). See also Supplemental Fig. S2D. Quantification of an experimental replicate is shown in Fig. S2E-F. (G) Plasmid DNA was replicated using *Xenopus* egg extracts in the presence of [α-^32^P]dATP to label newly-synthesized DNA strands under TOP2 inhibited (TOP2-i) conditions and immunodepleted of TOP3α, BLM, or both. (H) Replication was performed as in (G). Samples were separated on a native agarose gel and visualized by autoradiography. See also Supplemental Fig. S2G. (I) Quantification of fork merger from (H).

TOP2-i treatment traps TOP2 on DNA, which exacerbates fork merger defects and induces aberrant DNA species (HMws; Fig. 1B) (*6*). Thus, to more directly examine the relationship between TOP2α and TOP3α we repeated our analysis (Fig. 1A-D) in TOP2α-immunodepleted extracts (Fig. 1E) where TOP2-i was included to block residual TOP2 activity (*6*). TOP2α immunodepletion caused replication fork structures to persist (Fig. 1F, lanes 1-4) but to a lesser extent than following TOP2-i treatment (Fig. 1B, lanes 1-4), leading to efficient catenane formation (cats; Fig. 1F. lanes 1-4), as expected (*6*). Total DNA synthesis was unaffected (Fig. 1G), consistent with TOP2α immunodepletion interfering with termination and not earlier stages of DNA replication, as previously described (*6*). Importantly, TOP3α immunodepletion delayed fork merger and this was rescued by addition of recombinant TRR (Fig. 1F, lanes 5-12, Fig. 1H). Thus, TOP3α promotes fork merger in the absence of TOP2α, not just upon TOP2α inhibition. The catenanes formed in TOP2α-immunodepleted extracts included a subset of slower migrating species that contained fewer intertwines (Cats-; Fig. 1F, lanes 1-4), also as previously found (*6*). Importantly TOP3α immunodepletion blocked formation of these species and this condition yielded only the faster-migrating catenanes with more intertwines (Cats+; Fig. 1F, lanes 5-8). This change in catenane distribution was rescued upon re-addition of recombinant TRR (Fig. 1F, lanes 9-12), which demonstrated that TOP3α promoted unlinking of catenanes, either during or after DNA replication. Overall, these data indicate that TOP3α promotes fork merger and unlinking of daughter DNA duplexes in the absence of TOP2α activity.

### TOP3α promotes fork merger independently of RMI1-RMI2-BLM

Our initial analysis (Fig. 1) indicated that TOP3α or another component of the TRR complex promotes fork merger. Additionally, TOP3α can play a non-enzymatic structural role (*53, 54*) and can also form the BLM-TOP3α-RMI1-RMI2 (BTRR) complex (*55–57*). We therefore investigated which TOP3α functions and interacting partners were required to promote fork merger when TOP2α activity is compromised (as in Fig. 1).

We first tested the requirement for the catalytic activity of TOP3α. To this end we purified TRR complex containing the Y367F mutant of TOP3α (Fig. S2A), which eliminates the conserved catalytic tyrosine residue (*65*), then tested its ability to rescue fork merger defects (Fig. 2A). The Wild Type TRR complex rescued fork merger (WT; Fig. 2B lanes 5-8, Fig. 2C) and did not impact DNA synthesis (Fig. S2B), as expected (Fig. 1A-D). In contrast, the catalytic mutant was indistinguishable from the control (Mut; Fig. 2B lanes 9-12, Fig. 2C). Thus, the ability of the TRR complex to promote fork merger was entirely dependent on the catalytic activity of TOP3α.

We next tested whether TOP3α itself could promote fork merger, or whether RMI1-RMI2 was required. We first immunodepleted RMI1 but found that this co-depleted TOP3α (Fig. S2C), consistent with reports that TOP3α forms a stable complex with RMI1-RMI2. This prevented us from independently assessing the contributions of RMI1-RMI2 by immunodepletion. We therefore tested the ability of TOP3α alone to rescue fork merger defects, compared to the complete TRR complex. We were unable to obtain catalytically active *Xenopus* TOP3α independent of RMI1-RMI2. Instead we used human TOP3α (hTOP3α). Surprisingly, hTOP3α alone was able to promote fork merger to the same extent as the *Xenopus* TRR complex (Fig. 2D-F, Fig. S2E). This was not due to the ability of residual *Xenopus* RMI1-RMI2 in the extract because xRMI1 was co-depleted using an xRMI1 antibody for this experiment (as in Fig. S2C). Thus, TOP3α alone could promote fork merger. However, hTOP3α only partially rescued formation of catenanes with fewer intertwines (Cats-; Fig. 2E, Fig. S2D,F) suggesting RMI1-RMI2 play some role. Overall, these data show TOP3α alone can support fork merger.

Finally, we examined the role of the BLM helicase, which binds TOP3α-RMI1-RMI2 to form the BTRR complex. We used a previously validated antibody (*66*) to immunodeplete BLM from extracts (Fig. S2G). This antibody efficiently immunodepleted BLM and co-immunoprecipitated TOP3α (Fig. S2G lanes 5-7), but left most TOP3α in extracts (Fig. S2G lanes 1-4), indicating that most TOP3α was not bound to BLM. We then examined fork merger in extracts immunodepleted of TOP3α or BLM or both (Fig. 2G). Surprisingly, BLM immunodepletion did not appreciably impair fork merger in mock- or TOP3α-depleted extracts (Fig. 2H-I). Thus, BLM helicase did not appear to be involved in the fork merger role we had identified for TOP3α.

Altogether, these results (Fig. 2) indicate that the catalytic activity of TOP3α promotes fork merger, independent of RMI1-RMI2 and BLM helicase.

### TOP3α acts primarily during replication termination

We previously found that TOP2α unlinks DNA throughout replication (i.e. during elongation) to prevent buildup of topological stress that would otherwise stall termination (*6*). Similarly, TOP3α and its orthologs have been reported to operate throughout replication (*9, 33*) as well as during termination (*36*). We therefore investigated whether TOP3α operates throughout replication, during termination, or both (Fig. S3A).

To test when during replication TOP3α functions, we wanted to inhibit TOP3α at different stages of replication. However, no inhibitors of TOP3α were available. We therefore raised an antibody targeting an extended region of TOP3α and tested whether it could act as an inhibitor of TOP3α function. Addition of this TOP3α antibody had no effect on DNA replication when TOP2α was fully active (Fig. 3A, Fig. 3B, lanes 1-4, 9-12), as expected (Fig. S1D-H). However, upon addition of TOP2-i the TOP3α antibody blocked fork merger (Fig. 3B, lanes 5-8, 13-16), reproducing the immunodepletion effect (Fig. 1A-D). The antibody had no effect when TOP3α was absent (Fig. S3C-F), indicating that it was specific. Thus, the antibody generated acts as an inhibitory antibody that specifically targets TOP3α.

**Figure 3:**
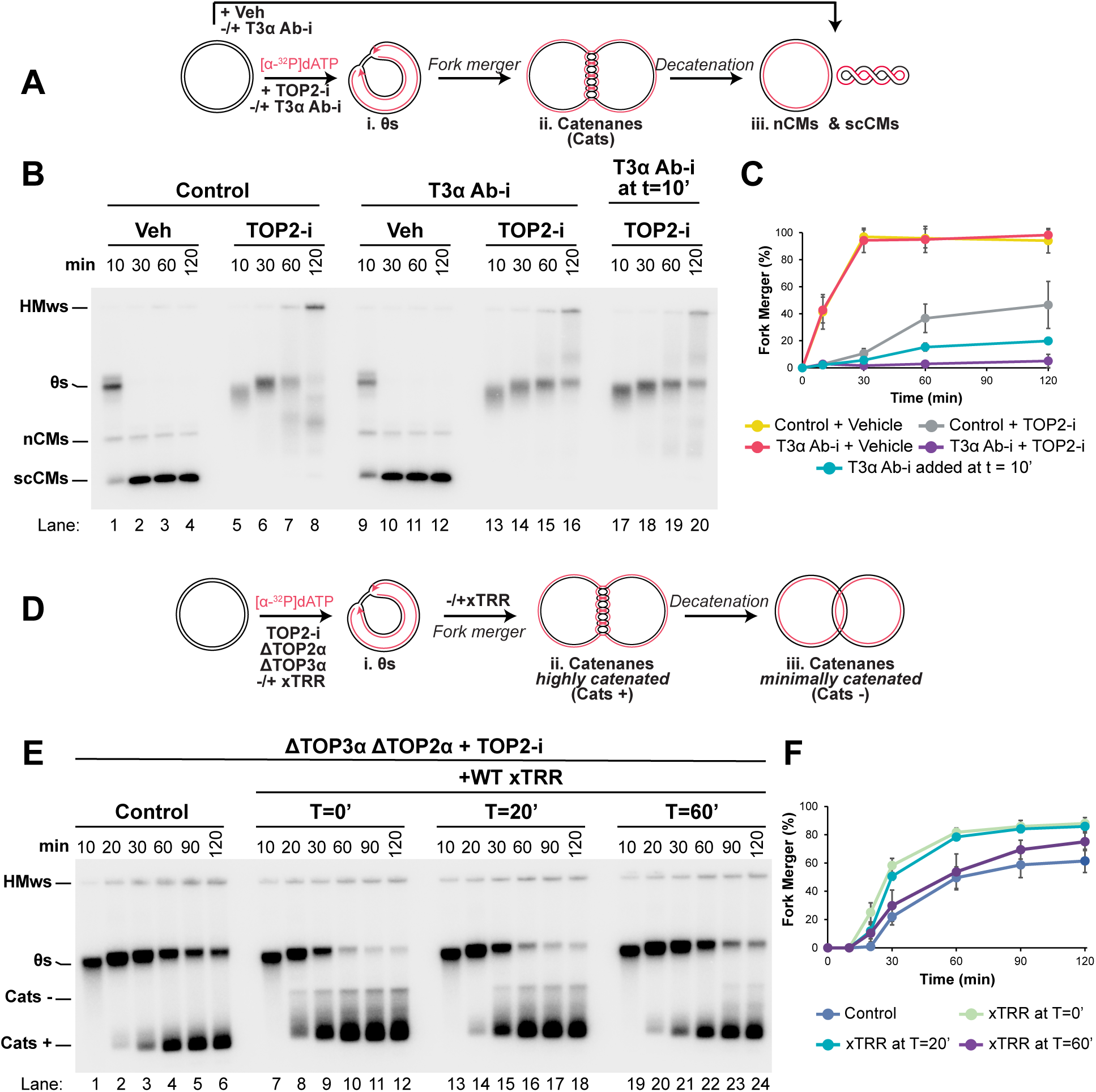
TOP3α acts on replication forks during termination. (A) Plasmid DNA was replicated using *Xenopus* egg extracts in the presence of [α-^32^P]dATP to label newly-synthesized DNA strands in the presence of vehicle (veh) or TOP2-i. At different time points extracts were supplemented with an inhibitory antibody targeting TOP3α (T3α Ab-i) or a control (Control). (B) Replication was performed as in (A). Samples were separated on a native agarose gel and visualized by autoradiography. (C) Quantification of fork merger from (B). See also Supplemental Fig. S3B. (D) Plasmid DNA was replicated using *Xenopus* egg extracts in the presence of [α-^32^P]dATP to label newly-synthesized DNA strands in extracts that were immunodepleted of TOP2α and TOP3α and supplemented with TOP2-i. At different time points recombinant xTRR was added back to the reaction. (E) Replication was performed as in (D). Samples were separated on a native agarose gel and visualized by autoradiography. See also Supplemental Fig. S3G-I. (F) Quantification of fork merger from (E).

To test whether TOP3α acts primarily late during replication we added TOP3α inhibitory antibody at 10 minutes, when most DNA synthesis was complete (∼70% complete; Fig. S3B). Antibody addition at this late time point reproduced most of the fork merger defect observed when the antibody was added at the onset of replication (Fig. 3B, lanes 5-8, 13-20, Fig. 3C). We note that this result likely underestimates the extent to which TOP3α functions during termination because the antibody may take time to fully inhibit TOP3α. Still, these data show that most of the effect of TOP3α on fork merger can be attributed to it functioning late during replication.

As an alternative approach to assess the role of TOP3α at different stages of DNA replication, we immunodepleted TOP2 and TOP3, then added back TRR either late during replication or at the onset of DNA replication (Fig. 3D). Analogously to the previous result (Fig. 3A-C), we found that addition of TRR rescued fork merger to the same extent, whether added at the onset of DNA replication (T=0’; Fig. 3E, lanes 7-12, Fig. 3F) or late during replication (T=20’; Fig. 3E, lanes 13-18, Fig. 3F). Although immunodepletion resulted in nonspecific slowing of DNA synthesis (compare Fig. S3B to Fig. S3G), this did not affect our experimental interpretation because we added xTRR after most DNA synthesis was complete (20 minutes; Fig. S3G). These data indicate that most of the effect of TOP3α on fork merger can be attributed to its role during termination.

Although fork merger was unaffected by the timing of TOP3α add back, there was a small difference in the number of less intertwined catenanes (Cats-; Fig. 3D) produced when TRR was added back late during replication (Fig. 3E, lanes 7-18, Fig. S3H-I). We therefore added back TRR even later during replication (T=60’; Fig. 3E, lanes 19-24) when most fork merger was complete (Fig. 3F). Doing so eliminated most of the less intertwined catenanes (Fig. S3H-I) even though the extent of replication was essentially unaltered between the two time points (Fig. S3G). Thus, once forks merge the ability of TOP3α to carry out unlinking is substantially diminished, suggesting that it is only able to act at forks and not on fully replicated DNA.

We previously showed that MCM10 and RTEL1 promote fork merger in the absence of TOP2α and this reflects a generalized role for these proteins in overcoming replication fork stalling (*67*). To determine whether TOP3α functions the same way, we assessed the impact of TOP3α immunodepletion on fork progression through an array of LacR proteins, which stalls replication forks (Fig. S3J). Immunodepletion of TOP3α had no effect on the ability of forks to progress through a LacR array (Fig. S3K-L). Thus, the role of TOP3α in promoting fork merger is specific to topological stress and does not reflect a general role for TOP3α in overcoming fork stalling.

Together, these data (Fig. 3) show that TOP3α operates primarily at replication forks during replication termination, rather than throughout DNA replication or as a general response to fork stalling.

### TOP3α targets lagging strand ssDNA

TOP3α may target lagging strand template ssDNA (‘lagging ssDNA’) (*24, 33*), which should be present throughout replication (Fig. S4Ai). However, it could equally target ssDNA generated by uncoupling (Fig. S4Aii) (*44–47*) or dsDNA given that it can unlink double-stranded DNA catenanes (Fig. S4Aiii) (*49, 50*). We therefore set out to directly test the model that TOP3α targets lagging strand ssDNA. The major prediction of this model is that increased lagging ssDNA should result in increased TOP3α activity. In our experiment (Fig. 1) this would mean that increased lagging strand ssDNA should at least partially rescue the decatenation defect in TOP2α mutants in a TOP3α-dependent manner (Fig. S4B). To test this, we took advantage of the Polα inhibitor CD437 (Polα-i) (*68*), which we found to specifically inhibit lagging strand synthesis in *Xenopus* egg extracts (Fig. S4C-D), as in cells (*69, 70*).

To test the effect of lagging ssDNA on catenane resolution, we replicated plasmid DNA in extracts immunodepleted for either TOP2α or both TOP2α and TOP3α, in the presence or absence of a partially inhibitory dose of Polα-i, and quantified the abundance of the less-catenated Cats- (Fig. 4A) (*6*) as a read-out for catenane abundance. We omitted TOP2-i from the reaction, which would have blocked formation of Cats- (*6*). Addition of Polα-i reduced the total amount of DNA synthesis (Fig. S4E), which is expected because Polα-Primase activity is required for initial priming of the leading strand during DNA replication (*71*). Strikingly, Polα-i treatment greatly increased the proportion of less intertwined catenanes formed (Fig. 4B, lanes 1-8, Fig. 4C, Fig. S4F) in TOP2α-immunodepleted extracts, which indicated that decatenation was increased by Polα-i treatment. Importantly, this effect was severely reduced in extracts immunodepleted of TOP2α and TOP3α (Fig. 4B, lanes 9-16, Fig. 4C, Fig. S4F). Thus, increased lagging ssDNA promotes decatenation in a manner that is dependent on TOP3α.

**Figure 4:**
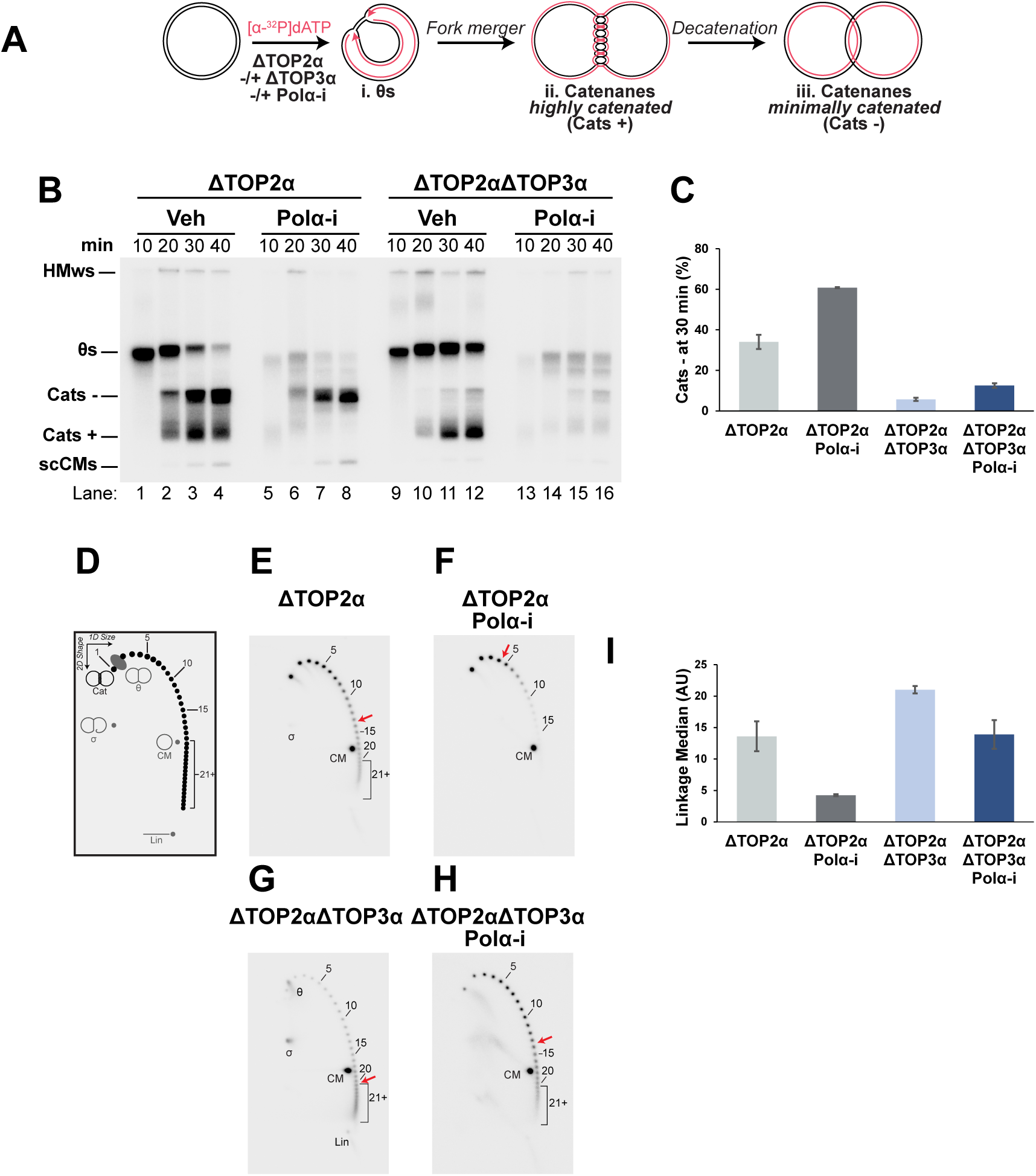
TOP3α acts on lagging strand template ssDNA. (A) Plasmid DNA was replicated using *Xenopus* egg extracts in the presence of [α-^32^P]dATP to label newly-synthesized DNA strands in extracts that were depleted of TOP2α with or without co-depletion of TOP3α. Extracts were also treated with either a small molecule inhibitor of POLα (POLα-i, CD437) or vehicle control (veh). (B) Replication was performed as in (A). Samples were separated on a native agarose gel and visualized by autoradiography. (C) Quantification of the lower mobility species of catenanes (Cats-) in (B). See also Fig. S4 E-F. (D) Schematic of the different species of catenated plasmid dimers detected by 2-D gel electrophoresis following nicking of plasmid DNA to relieve supercoiling. Each spot on the indicated arc corresponds to a species of catenanes with a distinct number of linkages. Species with 21 or more linkages cannot be discerned from each other because the individual positions are too close to be resolved. (E) The sample from Lane 3 in (B) was separated by 2-D gel electrophoresis and visualized by autoradiography. Note that only the section of the gel containing catenated species was analyzed. (F) The sample from lane 7 in (B) was analyzed as in (E). (G) The sample from lane 11 in (B) was analyzed as in (E). (H) The sample from lane 15 in (B) was analyzed as in (E). (I) Quantification of the median number of linkages for the catenated species detected in (E)-(H).

Formation of less intertwined catenanes (Cats-; Fig. 4A-C) is an indirect readout of decatenation. To directly examine decatenation we performed 2-D gel electrophoresis to measure the number of intertwines in the catenated products of DNA replication (Fig. 4D-I). We analyzed samples from the 30 minute time point of Fig. 4B when catenanes had initially formed (Fig. S4F) to prevent residual decatenation activity from remaining TOP2α from confounding the results. Following TOP2α immunodepletion the linkage median was approximately 14 (Fig. 4E, Fig. 4I). This was lower than we had previously reported (*6*), because omission of TOP2-i allowed some amount of unlinking to take place. Addition of Polα-i shifted the distribution of linkages down, resulting in a reduction to approximately 4 (Fig. 4F, Fig. 4I), consistent with our assessment that Polα-i reduces the number of intertwines in the catenated products of replication (Fig. 4A-C). Immunodepletion of TOP3α shifted the distribution of linkages up and the median was approximately 20 (Fig. 4G, Fig. 4I). Addition of Polα-i once again reduced the linkage median to approximately 14 (Fig. 4H, Fig. 4I) resulting in a smaller reduction than when TOP3α was present. There was no net effect on linkage median when TOP3α was immunodepleted and Polα-i was added, as evidenced by a linkage median of approximately 14 in both cases (Fig. 4E, Fig. 4H, Fig. 4I). Thus, increased lagging ssDNA reduces the number of intertwines in the catenated products of replication through a mechanism that involves TOP3α.

Collectively, our data (Fig. 4) provide direct evidence for the model that TOP3α acts on lagging ssDNA.

### TOP3α resolves intramolecular ssDNA linkages

Our finding that TOP3α specifically acts on lagging ssDNA raises the possibility that TOP3α may be uniquely important when lagging strand DNA synthesis is inhibited. To test this possibility, we replicated plasmid DNA in the presence of a high concentration of Polα-i using either mock- or TOP3α-immunodepleted extracts (Fig. 5A). No major differences in total DNA synthesis were observed (Fig. S5A). However, we observed the formation of a unique DNA species, that migrated slightly faster than nicked circular monomers, specifically in TOP3α-immunodepleted conditions (CM*; Fig. 5B, lanes 1-10, Fig. 5C). Thus, in the presence of elevated lagging ssDNA, TOP3α suppresses the formation of a unique DNA species.

**Figure 5:**
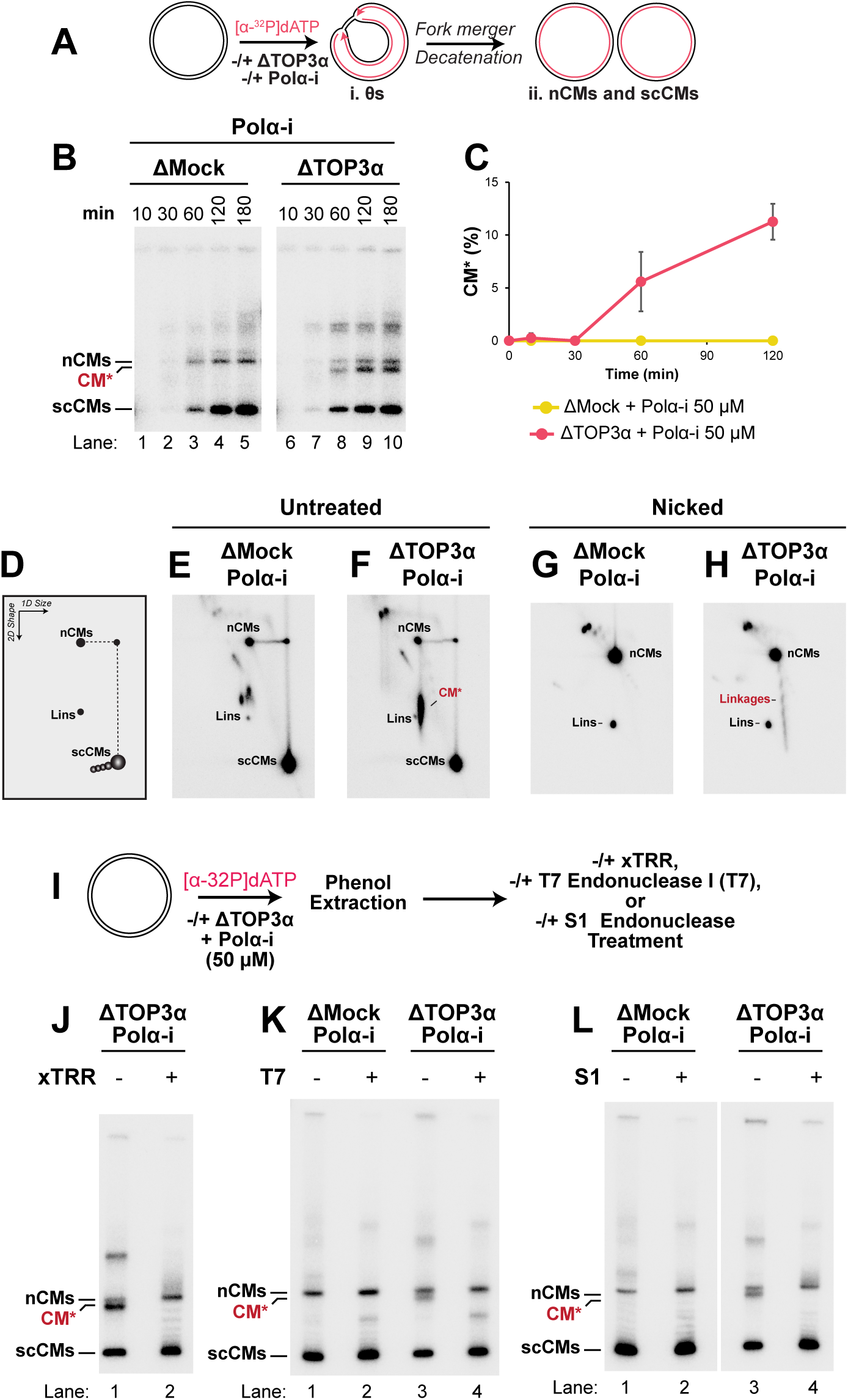
TOP3α resolves intramolecular ssDNA intertwines. (A) Plasmid DNA was replicated using *Xenopus* egg extracts in the presence of [α-^32^P]dATP to label newly-synthesized DNA strands in extracts that were treated with POLα-i with or without immunodepletion of TOP3α. (B) Replication was performed as in (A). Samples were separated on a native agarose gel and visualized by autoradiography. (C) Quantification of CM*, which is the circular monomer species migrating between supercoiled (scCMs) and nicked (nCMs) circular monomer species, in (B). See also Supplemental Fig. S5A. (D) Schematic of the migration of different monomeric species of circular monomers following 2-D gel electrophoresis. (E) The sample from lane 5 in (B) was separated by 2-D gel electrophoresis and visualized by autoradiography. Note that only the section of the gel containing monomeric species was analyzed. (F) The sample from lane 10 in (B) was analyzed as in (E). (G) The sample from lane 5 in (B) was nicked using a site-specific nicking enzyme then analyzed as in (E). (H) The sample from lane 10 in (B) was nicked using a site-specific nicking enzyme then analyzed as in (E). (I) Plasmid DNA was replicated using *Xenopus* egg extracts in the presence of [α-^32^P]dATP to label newly-synthesized DNA strands in extracts that were treated with POLα-i and mock- or TOP3α-immunodepleted. Replication products were then purified and treated with recombinant xTRR, T7 nuclease, or S1 nuclease. (J) Replication was performed as in (I). The purified products of replication in TOP3α-immunodepleted extracts were treated with either purified xTRR or buffer control then separated on a native agarose gel and visualized by autoradiography. Quantification is shown in Supplemental Fig. S5E-F. (K) Replication was performed as in (I). Purified replication products were treated with either T7 nuclease or buffer control then separated on a native agarose gel and visualized by autoradiography. Quantification is shown in Supplemental Fig. S5G-H. (L) Replication was performed as in (I). Purified replication products were treated with either S1 nuclease or buffer control then separated on a native agarose gel and visualized by autoradiography. Quantification is shown in Supplemental Fig. S5I-J.

To gain additional information about the structure of the unique DNA species we performed 2-D gel electrophoresis (Fig. 5D-F). The species migrated within the area corresponding to monomers and migrated similarly to linear species. However, when the samples were nicked with a site-specific nicking enzyme the species was predominantly converted to nicked circular monomer (Fig. 5G-H), indicating that it was a topoisomer of covalently closed circular monomer. We thus named the species CM* to reflect its identity as an altered Circular Monomeric plasmid.

We noticed that nicking CM* led to formation of an arc of topoisomers emanating from the nicked circular monomers (linkages; Fig. 5H). This pattern would typically be interpreted as intramolecular double-stranded DNA (dsDNA) knots. However, we disfavored this possibility because TOP3α appears to act on ssDNA during DNA replication (Fig. 3, Fig. 4), which made it difficult to envisage how its absence could lead to dsDNA knots. We reasoned that these species (linkages; Fig. 5H) could correspond to ssDNA linkages, in which case we would expect them to be resolved by nicking both DNA strands (i.e. by allowing both strands to freely rotate). Importantly, nicking both strands would not be expected to resolve dsDNA knots as breakage of both strands along with strand passage is required. Nicking both DNA strands did indeed resolve the linkages (Fig. S5B-C). Thus, the properties of CM* are consistent with the presence of some intramolecular ssDNA linkages.

The best characterized substrate for TOP3α is hemicatenanes (*22*), which are ssDNA linkages between different DNA molecules (Fig. S5D). We reasoned that the intramolecular ssDNA linkages present in CM* (Fig. 5H) should exhibit similar behavior (Fig. S5D). We therefore treated CM* with purified TRR complex (Fig. 5I-J), which resolves hemicatenanes. TRR was able to directly resolve CM* (Fig. 5J, Fig. S5E) while having minimal effect on supercoiled circular monomers (Fig. S5F), supporting the idea that it contained hemicatenane-like structures. We reasoned that any such structures should also be hypersensitive to ssDNA endonucleases so we also tested whether they could be resolved by T7 endonuclease (Fig. 5I, Fig. 5K, Fig. S5G-H) or S1 endonuclease (Fig. 5I, Fig. 5L, Fig. S5I-J). CM* was resolved by both T7 endonuclease (Fig. 5K, compare lanes 3-4) and S1 endonuclease (Fig. 5L, compare lanes 3-4). Under these same conditions, supercoiled plasmids were unaffected indicating that this effect was specific to single-stranded DNA (scCMs; Fig. 5K, lanes 1-2; Fig. S5H; Fig. 5L, lanes 1-2; Fig. S5J). Thus, the properties of CM* suggest it contains intramolecular single-stranded linkages (Fig. S5K).

Overall, our data (Fig. 5) show that high levels of lagging ssDNA create a requirement for TOP3α to suppress the formation of intramolecular ssDNA linkages.

## DISCUSSION

We have identified a specialized role for TOP3α during replication termination, where it can compensate for loss of TOP2α activity in order to promote fork merger (Fig. 6A-B). To do so, TOP3α targets lagging single-stranded DNA (ssDNA) independent of BLM and RMI1-RMI2. We also found that TOP3α plays an additional role in resolving intramolecular ssDNA intertwines that can arise in the presence of high levels of lagging ssDNA (Fig. 6C). Thus, TOP3α resolves both inter- and intra-molecular intertwines during vertebrate DNA replication.

**Figure 6:**
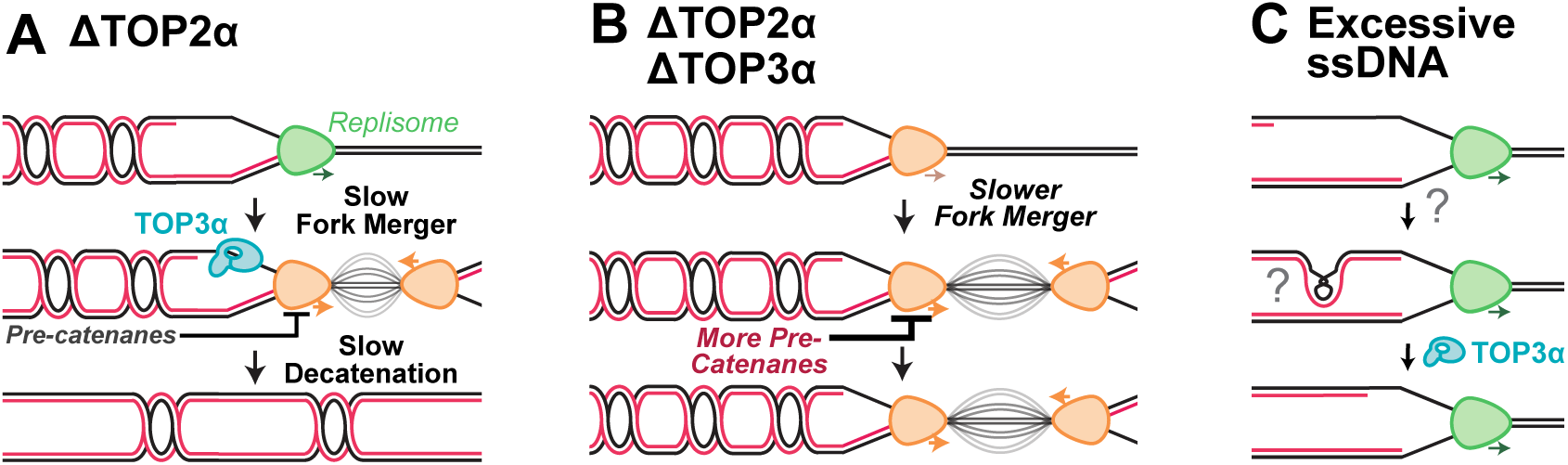
Model for distinct functions of TOP3α during vertebrate DNA replication. (A) In the absence of TOP2, pre-catenanes accumulate and cause converging replication forks to stall, leading to slowed fork merger. TOP3α resolves pre-catenanes by acting on lagging template ssDNA during termination, contributing to residual fork merger and decatenation. (B) In the absence of both TOP2 and TOP3 pre-catenanes persist, resulting in prolonged replication fork stalling. (C) In the presence of high levels of lagging ssDNA intramolecular ssDNA intertwines can form and these can be resolved by TOP3α.

### TOP3α function during DNA replication

Our finding that TOP3α exerts no detectable effect during normal DNA replication in *Xenopus* egg extracts (Fig. S1D-H) differs from a previous study that found a role for TOP3α (*38*). We favor the idea that the cellular role for TOP3α during replication reflects some facet of nuclear biology that is missing from the *Xenopus* egg extract system, such as transcription-replication collisions (*39*) given that the extracts we use are transcriptionally inactive unless provided with altered DNA template or additional transcription factors (*72, 73*). A role outside of unperturbed fork progression would be distinct from bacteria, where topo III acts throughout replication (*33*) and this difference may reflect more extensive lagging strand ssDNA at bacterial replication forks.

We previously reported that TOP2α resolves pre-catenanes and loss of TOP2α causes replication forks to stall during termination (*6*). Here we find that TOP3α similarly resolves pre-catenanes when TOP2α is absent and loss of TOP3α exacerbates stalling during termination (Fig. 1). In contrast to TOP2α, which resolves pre-catenanes throughout replication (*6*), TOP3α predominantly resolves pre-catenanes during termination (Fig. 3). We favor a model whereby TOP2α resolves most pre-catenanes and thus obviates a need for TOP3α under normal conditions, but when TOP2α function is absent TOP3α can act at termination to resolve any remaining pre-catenanes.

We previously found that the helicase RTEL1, recruited by the replisome component MCM10, promotes fork merger during termination (*67*). As for TOP3α, MCM10-RTEL1 is crucial for fork merger when forks stall due to lack of TOP2α. However, MCM10-RTEL1 promotes fork progression as part of a generalized response to stalling that is not specific to topological stress or termination (*67*). In contrast, we found that TOP3α acts when forks stall during termination due to topological stress (Fig. 3) and not when forks are stalled at a replication barrier (Fig. S3J-L). Thus, MCM10-RTEL1 and TOP3α can both support fork progression when forks stall during termination but operate through distinct mechanisms.

Loss of both TOP2α and TOP3α leads to a synergistic block to fork convergence (Fig. 1). In human cells, a similar synergistic effect on accumulation of ultrafine anaphase bridges occurs when activity of both TOP2α and TOP3α is inhibited (*58*). Importantly, anaphase bridges can arise from unreplicated stretches of DNA (*74*) that could reflect termination defects. Thus, the role we have identified for TOP3α may contribute to its role in suppressing anaphase bridges, which is critically important for mitosis.

### TOP3α Independence from BLM and RMI1-RMI2

We identified a nuclear role for TOP3α that is independent of BLM and RMI1-RMI2 (Fig. 2). Fork merger by TOP3α was unaltered by the loss of BLM and TOP3α was sufficient to rescue the TOP3α immunodepletion effect. However, BLM and RMI1-RMI2 are clearly important for TOP3α function, so this observation suggests that some facet of TOP3α unlinking at replication forks obviates the requirement. For example, the replicative CMG helicase may substitute for the unwinding activity of the BLM helicase and other protein-protein interactions may substitute for RMI1-RMI2 function. We note that the independence of RMI1-RMI2-BLM matches the situation during mitochondrial DNA replication (*40*), suggesting it could be generally true of TOP3α function during DNA replication. Given that BLM and RMI1-RMI2 play crucial roles in cells (*56, 75–77*) and during other aspects of replication (*66*) it will be important to determine in which circumstances they are required for TOP3α activity.

### TOP3α acts on Lagging Strands

Our data show that impaired lagging strand synthesis increased daughter strand unlinking and this increase was dependent on TOP3α (Fig. 4). Thus, we provide direct evidence that TOP3α targets lagging ssDNA. We found no ability of TOP3α to resolve double-stranded DNA (dsDNA) intertwines, despite its ability to do so in a range of circumstances (*32, 49, 50*). These data argue that under normal circumstances TOP3α acts specifically on ssDNA structures to resolve DNA intertwines.

A key question is why TOP3α acts preferentially during termination even though lagging-strand ssDNA is present throughout replication. One possibility is that TOP3α requires persistent ssDNA but during fork progression lagging-strand ssDNA is too transient to be acted upon and persists long enough only when forks stall during termination. Alternatively, TOP3α may not be able to act on pre-catenanes during earlier stages of replication if they preferentially partition into the fully double-stranded regions of the daughter strands. Distinguishing these models will be an important goal of future work.

### TOP3α acts on intramolecular Intertwines

An unexpected outcome of our investigations was the discovery that high levels of lagging-strand ssDNA can give rise to intramolecular ssDNA intertwines (Fig. 5, Fig. 6C). To our knowledge, such structures have not been previously described in the context of DNA replication, though a potentially similar structure was speculated to arise from DNA looping (*78*). The simplest explanation for these structures is that they are essentially intramolecular hemicatenanes, which typically arise from Holliday junctions formed during HR (*22*). However, we disfavor this as a source of intramolecular ssDNA linkages because homologous recombination would be expected to favor sister molecules, whereas we instead observe a preference for formation of these intertwines within the same molecule. Thus, the origin of these structures is highly uncertain. Although these events are rare, they could conceivably give rise to highly mutagenic extrachromosomal DNA (*79*). An important future direction is to establish the exact causes and consequences of these intramolecular intertwines.

## Methods

### *Xenopus* Egg Extracts

*Xenopus* egg extracts were prepared from *Xenopus laevis* wild-type males and females (Nasco) as previously described (*80*). All experiments and protocols were approved by the Vanderbilt University Division of Animal Care (DAC) and Institutional Animal Care and Use committee (IACUC).

### Plasmid Construction and Preparation

The construction of pJD152 (p[lacOx16]) and pJD156 (p[lacOx32]) was described previously (*81*). pJD156 (p[lacOx32]) was used in Fig. S3J-L. All other experiments used pJD152(p[lacOx16]) as a template for replication.

### Protein Purification

Biotinylated LacR protein was expressed in *Escherichia coli* bacteria cells and purified as described previously (*61, 81*).

Recombinant wild-type (WT) and catalytic mutant (TOP3α Y367F mutant) *Xenopus* TOP3α-RMI1-RMI2 complex (xTRR) were expressed and purified by Genscript (U1353074G0 for WT; U062D412G0 for mutant). xTRR was expressed in insect cells as a complex, then purified via affinity purification against HIS and FLAG tags. In Fig. 1 WT xTRR was used at a final concentration of 42.68 µg/ml. In Fig. 2 wild type xTRR was used at a final concentration of 5.12 µg/ml while mutant xTRR was used at a final concentration of 13.35 µg/ml. These concentrations of WT and mutant xTRR were selected quantitatively for having the same amount of TOP3α protein (see Fig. S2A).

To purify human TOP3α, Rosetta2 (DE3) pLysS E. coli cells (Novagen, Sigma) were transformed with the pET29H2 plasmid (*82*), which encodes N-terminally His-tagged hTOP3A, and grown in 2YT media supplemented with kanamycin (50 µg/ml) and chloramphenicol (20 µg/ml) to OD600 of 0.5. Cells were cooled down and IPTG was added to a final concentration of 1 mM. Expression was carried out overnight at 18°C with constant shaking. Cells were harvested by centrifugation, resuspended in lysis buffer (20 mM Hepes-NaOH pH = 7.5, 1 M NaCl, 0.5 mM TCEP, 10% v/v glycerol, 0.1% v/v Triton X-100, 30 mM imidazole, 1 mM PMSF, and Roche Complete EDTA-free protease inhibitor), and lysed by sonication. After removing cell debris by centrifugation (Sorvall SS34 rotor, 18k rpm, 40 min), the soluble fraction was loaded onto a 5 ml Nickel-NTA column (HisPur Ni-NTA resin, Thermo Fisher) equilibrated with lysis buffer. After loading, the column was washed with lysis buffer and a wash buffer (20 mM Hepes-NaOH pH = 7.5, 0.2 M NaCl, 0.5 mM TCEP), and the protein was eluted in wash buffer supplemented with 250 mM imidazole. The eluted sample was diluted 1:1 with wash buffer and loaded onto a 5 ml HiTrap Heparin HP (Cytiva) column equilibrated with the same buffer. Protein was eluted with a NaCl gradient of 0.2 M to 1 M in wash buffer. Elution fractions were run on an SDS-PAGE gel and samples containing TOP3A were pooled. The pooled sample was concentrated to ∼1 mg/ml with a 50 kDa ultra centrifugation filter (Amicon) and dialyzed into storage buffer (20 mM Hepes-NaOH pH = 7.5, 0.2 M NaCl, 10% v/v glycerol, 1 mM DTT). Concentration was determined using UV absorbance and aliquots were frozen in an ethanol/dry ice bath and stored at −80°C.

### Antibodies

For immunodepletions, a TOP3α antibody (NEP 3858) was generated in rabbits by New England Peptide using purified peptide of *Xenopus* TOP3α composed of the sequence CHQPGHNRTNSPQNR.

For Western Blotting and inhibition of TOP3α a second TOP3α antibody was generated. The following HIS-tagged polypeptide composed of the sequence MHHHHHHHHHHNRPPQAAPRESNHLQVNQTSNRMGNSGDGRGRQPQTKTLKPPVVKPSTSRGAPPIPGNSESNEVVCNCGVTAVQLTVRKEGPNQGRPFYKCNGGACNFFLWADQEDGGGQNAPAGPSTGRPSNAFNNSRAPDSFRSGGGSNDPGSEGQAVVCMCNQPAVTRTVQKDGANKGRQFHTCSKPREQQCGYFQWADENVPPGSSSGFSGSSGGSSFSGGSGAQRNSGKKNKRSENSFSGPTAKKPRSCG was purified by Genscript (U8068HWUG0). This antigen was then used to generate affinity purified TOP3α antibody (NEP 5606) in rabbits by New England Peptide.

*Xenopus* antibodies targeting TOP2α and BLM were previously generated (*6, 66*). RMI1 antibody was from Proteintech (Cat#: 14630-1-AP).

### Immunodepletions

To deplete TOP2α, TOP3α, BLM, and RMI1 from *Xenopus* egg extracts, 1.29 volumes Protein A-coupled magnetic beads (Dynabeads Protein A (30 mg/mL)) were bound to target antibody or control anti-KLH (Keyhole limpet hemocyanin) ‘mock’ antibody (0.5 μg antibody per 1 μl beads). Antibody-bound beads were then incubated with 1 volume of NPE or 0.5 volumes HSS for 20 min at room temperature with end-over-end rotation and this was repeated once. Depleted extracts were then collected and used for DNA replication (as above). For co-depletion of two proteins, beads were combined for each round of depletion and the number of depletion rounds remained constant. In all experiments beads bound to control anti-KLH were used to ensure that the concentration of beads and anti-KLH was identical for all conditions.

### DNA Replication in *Xenopus* egg extracts

DNA replication was performed as previously described (*61*). High Speed Supernatant (HSS) was incubated with nocodazole (3 ng/μl) and ATP regenerating system(‘ARS,’ 20 mM phosphocreatine, 2 mM ATP and 5 ng/μl phosphokinase) for five minutes at room temperature. When indicated, LacR protein was bound to *lacO* repeats on plasmid DNA for one hour at room temperature before licensing DNA. To license plasmid DNA, one volume of ‘licensing mix’ was prepared by incubating plasmid DNA with HSS (final concentration: 6.75-13.5 ng/μl DNA) at room temperature for 30 minutes. NucleoPlasmic Extract (NPE) was supplemented with ARS, and DTT (final concentration: 2 mM). To initiate replication, 2 volumes of NPE mix were added to 1 volume of licensing mix. If applicable, replication forks that were stalled at a LacR-bound *lacO* array were released by IPTG addition as described previously (*61, 81*). Samples were withdrawn into Replication Stop Solution (8 mM EDTA, 0.13% phosphoric acid, 10% ficoll, 5% SDS, 0.2% bromophenol blue, 80 mM Tris, pH 8) then vigorously mixed and treated with Proteinase K (929 ng/μl). Polymerase α inhibitor (Polα-i, CD437) was dissolved in DMSO and used at a final concentration of 6 µM for Fig. 4, 200 µM for Fig. S4C-D, and 50 µM for Fig. 5. Polα-i was used with a final concentration of 4% (v/v) DMSO in the reaction.

### Analysis of replication intermediates

Replication intermediates were separated by native agarose gel electrophoresis and detected by autoradiography. Quantification was performed using ImageJ (NIH). To quantify total DNA synthesis, total lane signal was measured and normalized to the maximum signal across all time points and conditions (AU). The abundance of supercoiled monomers, linears, and other indicated species are expressed as a percentage of total lane signal (%).

### 2-D Gel Electrophoresis

Nicked 2-D catenane gels (Fig. 4) were performed as described previously (*6*). Briefly, purified DNA was digested with 0.02 U/µL Nb.BtsI (New England Biolabs) and then separated on a 0.4% agarose gel at 1 V/cm for 21.5 hours. The gel was stained with 0.3 mg/ml ethidium bromide then lanes were excised and recast in a 1.2% gel containing 0.3 mg/ml ethidium bromide before being separated at 4.8 V/cm for 12 hours. Radiolabeled DNA was detected by phosphorimaging. Total catenane signal was quantified as a percentage of signal from the entire 2D gel autoradiograph. To calculate linkage median, the signal from every 5 catenanes (e.g., species with 1-5 linkages, species with 6-10 linkages, etc.) was measured and then expressed as a percentage (%) of total catenanes signal.

For Fig. 5, purified DNA was either left uncut or digested with 0.4 U/µL Nb.BssSI (New England Biolabs) to cut one strand only or 0.2 U/µL Nb.BtsI (New England Biolabs) to cut both strands. Digests were performed in rCutsmart buffer for 1 hour at 37°C. DNA was then separated on a 0.4% agarose gel at 1 V/cm for 21.5 hours. Lanes were excised and recast in a 1.2% gel containing 0.3 mg/ml ethidium bromide before being separated at 4.5 V/cm for 12 hours. Radiolabelled DNA was detected by phosphorimaging.

### DNA Enzyme Digests

For all DNA enzyme digests, DNA samples were first purified via phenol-chloroform extraction and then precipitated by ethanol precipitation before resuspension in 10mM Tris (pH 8.0) buffer. For S1 Nuclease (Thermo Scientific, Cat#: EN0321) digestion, S1 nuclease was diluted with 1x S1 Nuclease Reaction Buffer (Thermo Scientific Cat#: EN0321) and then approx. 10 ng of DNA was treated with 1 Unit S1 Nuclease (Fig. 5) in 1x S1 Nuclease Reaction Buffer at 23°C for 30 minutes. Reactions were stopped by addition of 1 µl of 0.5 M EDTA and heated at 70°C for 10 minutes, as per manufacturer’s instructions. 3 µl of 6x SDS Loading DNA Buffer (15% Ficoll-400, 66.5 mM EDTA, 20 mM Tris–HCl pH 8.0, 0.1% SDS, 0.09% Bromophenol blue) was added to each sample reaction before loading onto a 1% agarose TBE native gel. DNA products were separated at 5 V/cm for approximately 2 hours.

For T7 Endonuclease I (New England Biolabs, Cat#: M0302S) treatment, approximately 10 ng of DNA was treated with 3 U/ml T7 Endonuclease I (Fig. 5) in 1x Buffer 2 (New England Biolabs) at 23°C for 30 minutes. Reactions were stopped by addition of 2 µl of 6x SDS Loading DNA Buffer (15% Ficoll-400, 66.5 mM EDTA, 20 mM Tris–HCl pH 8.0, 0.1% SDS, 0.09% Bromophenol blue). DNA was loaded onto a 1% agarose TBE native gel and products were separated at 5 V/cm for approximately 2 hours.

### Topoisomerase Treatment

For *Xenopus* Topoisomerase IIIα-RMI1-RMI2 (xTRR) (Genscript) protein treatment, approximately 10 ng of DNA was treated with a final concentration of 2.24 μg/ml xTRR protein at 37°C for 15 minutes in a buffer containing 35 mM Tris-HCl, pH 7.5, 3 mM MgCl2, 60 mM NaCl, 20% glycerol, 1 mM dithiothreitol (DTT), 2 mM ATP, and 20 mM phosphocreatine. 1 unit of Creatine Phosphokinase was added last to each reaction. During the incubation, TopSTOP buffer was prepared from 20 μL Proteinase K (929 ng/μl), 80 μL filtered MilliQ water, and 100 μL 6% SDS. After incubation, a volume of TopSTOP buffer equivalent to a third of the reaction was added and then the reaction was incubated for a further 15 minutes at 37°C. Reactions were stopped by addition of 3 µl of 6x SDS Loading DNA Buffer (15% Ficoll-400, 66.5 mM EDTA, 20 mM Tris–HCl pH 8.0, 0.1% SDS, 0.09% Bromophenol blue). DNA was loaded onto a 1% agarose TBE native gel and products were separated at 5 V/cm for approximately 2 hours.

## Acknowledgements

JMD was supported by R35GM128696. WGD was supported by R01GM043974. KCN and GMH are supported by the Intramural Research Program of the National Heart, Lung, and Blood Institute (NHLBI), National Institutes of Health (NIH) (1ZIAHL001056 to K.C.N.). The contributions of the NIH authors are considered Works of the United States Government. The findings and conclusions presented in this paper are those of the author(s) and do not necessarily reflect the views of the NIH or the U.S. Department of Health and Human Services.

## Author Contributions

JMD conceptualized and supervised the project and wrote the manuscript. SVR performed all experimental work and analysis except where otherwise noted. LVC made the initial discovery of synergy between inactivation of TOP2α and TOP3α. GMH purified human TOP3α with supervision from KCN. DRH performed the experiment in Fig. S4C-D. SND performed the experiment in Fig. S3F. WGD supplied BLM antibody.

## Supplemental Figure Legends

**Supplemental Figure S1:** Roles of TOP3α during vertebrate DNA replication

(A) Cartoon depicting the different stages of vertebrate DNA replication during which TOP3 could act: elongation (i) and the fork merger stage of termination (ii) at lagging strand template ssDNA; and (iii) the decatenation stage of termination by acting on dsDNA catenanes.

(B) Cartoon depicting the role of TOP2 during vertebrate DNA replication termination. Under unperturbed conditions TOP2 primarily resolves catenanes behind replication forks to promote fork merger.

(C) Cartoon depicting a potential role for TOP3 during vertebrate DNA replication termination. In the absence of TOP2 catenanes accumulate, causing converging replication forks to stall. However, forks ultimately overcome stalling and limited decatenation takes place, which could in part reflect the activity of TOP3.

(D) Plasmid DNA was replicated using *Xenopus* egg extracts in the presence of [α-^32^P]dATP to label newly-synthesized DNA strands with or without TOP3α immunodepletion.

(E) TOP3α-immunodepleted extracts from (D) were analyzed by western blotting for TOP3α alongside a dilution series of mock-immunodepleted extracts. TOP3α is visible in the 100-fold dilution of mock-immunodepleted extract (lane 6) but not in the undiluted TOP3α-immunodepleted extract, suggesting >99% of TOP3α was removed.

(F) Replication was performed as in (D). Samples were separated on a native agarose gel and visualized by autoradiography.

(G) Quantification of total DNA synthesis from (F). Because DNA replication is asynchronous in this system, DNA synthesis reflects the rate of both initiation and elongation. No detectable difference was observed between mock- and TOP3α-immunodepleted extracts, indicating that TOP3α does not play a major role in initiation or elongation.

(H) The fork merger stage of termination was quantified from (F). No detectable difference was observed between mock- and TOP3α-immunodepleted extracts, indicating that TOP3α does not play a major role in fork merger.

(I) Purified *Xenopus* TOP3α-RMI1-RMI2 complex was analyzed by SDS-PAGE to assess purity. Three distinct bands are visible, corresponding to the different complex subunits and no other bands are discernable, indicating a high level of purity of the prep.

**Supplemental Figure S2:** Analysis of TOP3α-interacting proteins

(A) Purified wild type (WT) and catalytic mutant (Mut) of *Xenopus* TOP3α-RMI1-RMI2 complex were analyzed by SDS-PAGE to assess evenness of loading. Even levels of TOP3α are detected for both conditions (∼120 kDa band) but the mutant preparation contains higher levels of the RMI1 and RMI2 components (∼80 kDa and ∼16 kDa bands).

(B) Quantification of total DNA synthesis from Fig. 2B.

(C) Extracts were immunodepleted of RMI1 and then analyzed by western blotting for RMI1 and TOP3α. Arrows indicate expected band positions. RMI1 immunodepletion removed >95% of both RMI1 and TOP3α (compare lanes 1 and 5) indicating that TOP3α co-depletes with RMI1.

(D) Quantification of Cats- species from Figure 2E.

(E) Quantification of an experimental replicate of Figure 2F.

(F) Quantification of an experimental replicate of (D).

(G) Mock- and BLM-immunodepleted Extracts from Figure 2G were analyzed by western blotting for BLM and TOP3α. Arrows indicate expected band positions. * indicates background band adjacent to BLM that was not immunodepleted.

**Supplemental Figure S3:** Analysis of TOP3α activity during different stages of DNA replication

(A) Cartoon model depicting different stages of vertebrate DNA replication during which TOP3α acts.

(B) Quantification of total DNA synthesis from Fig. 3B.

(C) To assess the ability of a putative TOP3α inhibitory antibody (T3α Ab-i) to specifically inhibit TOP3α, plasmid DNA was replicated using *Xenopus* egg extracts in the presence of [α-^32^P]dATP to label newly-synthesized DNA strands under TOP2-immunodepleted and inhibited (TOP2-i) conditions, with or without TOP3α immunodepletion and addition of T3α Ab-i. As a control, antibody targeting keyhole limpet hemocyanin (Ctrl Ab-i) was used.

(D) Samples from (C) were separated on a native agarose gel and visualized by autoradiography. T3α Ab-i reproduced the fork merger defect of T3α immunodepletion (compare lanes 5-8 to 9-12) and there was no effect of T3α Ab-i when T3α was immunodepleted (compare lanes 9-12 to 13-16). Thus, T3α Ab-i specifically inhibits TOP3α.

(E) Fork merger was quantified from (D).

(F) Experimental replicate of (E).

(G) Quantification of total DNA synthesis from Fig. 3E.

(H) Quantification of the lower mobility species of catenanes (Cats-) in Fig. 3E.

(I) Bar Chart of the 120 minute time point from (H).

(J) To assess the role of TOP3α during replication fork stalling, plasmid DNA containing a *lacO* array was bound with LacR then was replicated using *Xenopus* egg extracts in the presence of [α-^32^P]dATP to label newly-synthesized DNA strands with or without TOP3α immunodepletion. Upon encountering the LacR barrier forks stall and then gradually progress through the *lacO* array until the two converging forks encounter each other and undergo fork merger, which acts as a read out for fork progression following stalling. Residual LacR binding interferes with decatenation and creates a range of complex topoisomers so to directly monitor fork merger, DNA was purified then restriction digested to monitor the conversion of high molecular weight replication fork structures (Double Ys) to low molecular weight linear molecules (linears).

(K) Samples from (J) were separated on a native agarose gel and visualized by autoradiography.

(L) Fork merger was quantified from (K). There was no difference in the progression of stalled forks between mock- and TOP3α-immunodepleted extracts, indicating that TOP3α does not play a general role in promoting stalled fork progression.

**Supplemental Figure S4:** Analysis of TOP3α activity on different replication fork structures

(A) Cartoon depicting the different replication fork structures TOP3α could act upon: lagging strand template ssDNA at replication forks (i); template ssDNA on both leading and lagging strands exposed by replication fork uncoupling (ii); and duplex DNA behind the replication fork (iii).

(B) Cartoon depicting the experimental test from Figure 4 of the model that TOP3α acts on lagging template ssDNA to resolve catenanes. Loss of TOP2 (ΔTOP2) results in accumulation of catenanes due to insufficient TOP2 activity. If TOP3α acts on lagging ssDNA template to resolve catenanes then increasing the amount of lagging ssDNA template by inhibition of Polα-Primase would be expected to reduce the number of catenanes and this reduction would be expected to depend on the presence of TOP3α.

(C) Cartoon depicting the experimental setup used to test the specificity of (Polα-i, CD437) for inhibition of lagging strand synthesis. Plasmid DNA containing a *lacO* array was bound with LacR then was replicated using *Xenopus* egg extracts in the presence of [α-^32^P]dATP to label newly-synthesized DNA strands. Once forks were stalled at the array, they were restarted by addition of IPTG along with either vehicle control or Polα-i.

(D) Replication was performed as in (C). At different time points, replication intermediates from different time points were purified, restriction digested, then analyzed by denaturing alkaline gel electrophoresis to monitor the nascent DNA strands from the leftward moving fork in (C). Prior to IPTG addition a single population of nascent DNA strands existed at a specific location, corresponding to forks stalled at the LacR barrier (lanes 1, 8), as expected. Following IPTG and Vehicle addition nascent strands increased in size as forks restarted (lanes 2-4) then were converted to high molecular weight ligated products once leading strands were ligated to downstream lagging strands (lanes 4-7). Following IPTG and Polα-i addition two populations of nascent strands were observed: a persistent stalled species that did not increase in size, corresponding to the expected stalled lagging strands (stalled lag; lanes 9-11); and a slower-migrating species that increased in size, corresponding to the expected restarted leading strands (restarted lead; lanes 9-11). Both species quickly resolved and were converted to ligated products (lanes 11-14) as expected if leading strand synthesis proceeded unhindered until encountering the downstream lagging strands then triggered ligation as normal. Thus, Polα-i results in stalling of a subset of nascent strands while leaving another subset unaffected, as expected for inhibition of lagging but not leading strand DNA synthesis.

(E) Quantification of total DNA synthesis from Figure 4B

(F) Quantification of Cats- from Figure 4B.

**Supplemental Figure S5: Characterization of** TOP3α function during replication in the presence of high levels of lagging ssDNA

(A) Quantification of total DNA synthesis from Fig. 5B.

(B) The sample from lane 5 in Fig. 5B was nicked on both strands using a site-specific nicking enzyme then analyzed as in Fig. 5E.

(C) The sample from lane 10 in Fig. 5B was nicked on both strands using a site-specific nicking enzyme then analyzed as in Fig. 5E.

(D) Cartoon depicting the composition of intramolecular ssDNA intertwines that could explain the results in Fig. 5A-H.

(E) Quantification of CM* from Fig. 5J.

(F) Quantification of scCMs from Fig. 5J.

(G) Quantification of CM* from Fig. 5K.

(H) Quantification of scCMs from Fig. 5K.

(I) Quantification of CM* from Fig. 5L.

(J) Quantification of scCMs from Fig. 5L.

(K) Depiction of possible structure of CM* species from Fig. 5

## References

1. J. J. Champoux, DNA topoisomerases: structure, function, and mechanism. Annu Rev Biochem 70, 369–413 (2001).

2. Y. Pommier, Y. Sun, S. N. Huang, J. L. Nitiss, Roles of eukaryotic topoisomerases in transcription, replication and genomic stability. Nat Rev Mol Cell Biol 17, 703–721 (2016).

3. A. Keszthelyi, N. E. Minchell, J. Baxter, The Causes and Consequences of Topological Stress during DNA Replication. Genes (Basel) 7, (2016).

4. J. M. Dewar, How Topoisomerases Relax and Disentangle the Genome During DNA Replication. SSRN (Preprint), (2026).

5. R. Bermejo, Y. Doksani, T. Capra, Y. M. Katou, H. Tanaka, K. Shirahige, M. Foiani, Top1-and Top2-mediated topological transitions at replication forks ensure fork progression and stability and prevent DNA damage checkpoint activation. Genes Dev 21, 1921–1936 (2007).

6. D. R. Heintzman, L. V. Campos, J. A. W. Byl, N. Osheroff, J. M. Dewar, Topoisomerase II Is Crucial for Fork Convergence during Vertebrate Replication Termination. Cell Rep 29, 422–436.e425 (2019).

7. J. T. Yeeles, T. D. Deegan, A. Janska, A. Early, J. F. Diffley, Regulated eukaryotic DNA replication origin firing with purified proteins. Nature 519, 431–435 (2015).

8. S. J. Brill, S. DiNardo, K. Voelkel-Meiman, R. Sternglanz, Need for DNA topoisomerase activity as a swivel for DNA replication for transcription of ribosomal RNA. Nature 326, 414–416 (1987).

9. H. Hiasa, K. J. Marians, Topoisomerase III, but not topoisomerase I, can support nascent chain elongation during theta-type DNA replication. J Biol Chem 269, 32655–32659 (1994).

10. H. Hiasa, K. J. Marians, Topoisomerase IV can support oriC DNA replication in vitro. J Biol Chem 269, 16371–16375 (1994).

11. H. Hiasa, K. J. Marians, Two distinct modes of strand unlinking during theta-type DNA replication. J Biol Chem 271, 21529–21535 (1996).

12. J. S. Minden, K. J. Marians, Replication of pBR322 DNA in vitro with purified proteins. Requirement for topoisomerase I in the maintenance of template specificity. J Biol Chem 260, 9316–9325 (1985).

13. L. Yang, M. S. Wold, J. J. Li, T. J. Kelly, L. F. Liu, Roles of DNA topoisomerases in simian virus 40 DNA replication in vitro. Proc Natl Acad Sci U S A 84, 950–954 (1987).

14. Y. Ishimi, K. Sugasawa, F. Hanaoka, T. Eki, J. Hurwitz, Topoisomerase II plays an essential role as a swivelase in the late stage of SV40 chromosome replication in vitro. J Biol Chem 267, 462–466 (1992).

15. D. H. Weinberg, K. L. Collins, P. Simancek, A. Russo, M. S. Wold, D. M. Virshup, T. J. Kelly, Reconstitution of simian virus 40 DNA replication with purified proteins. Proc Natl Acad Sci U S A 87, 8692–8696 (1990).

16. B. J. Peter, C. Ullsperger, H. Hiasa, K. J. Marians, N. R. Cozzarelli, The structure of supercoiled intermediates in DNA replication. Cell 94, 819–827 (1998).

17. J. Cebrián, A. Castán, V. Martínez, M. J. Kadomatsu-Hermosa, C. Parra, M. J. Fernández-Nestosa, C. Schaerer, P. Hernández, D. B. Krimer, J. B. Schvartzman, Direct Evidence for the Formation of Precatenanes during DNA Replication. J Biol Chem 290, 13725–13735 (2015).

18. S. DiNardo, K. Voelkel, R. Sternglanz, DNA topoisomerase II mutant of Saccharomyces cerevisiae: topoisomerase II is required for segregation of daughter molecules at the termination of DNA replication. Proc Natl Acad Sci U S A 81, 2616–2620 (1984).

19. J. J. Champoux, R. Dulbecco, An activity from mammalian cells that untwists superhelical DNA--a possible swivel for DNA replication (polyoma-ethidium bromide-mouse-embryo cells-dye binding assay). Proc Natl Acad Sci U S A 69, 143–146 (1972).

20. J. C. Wang, Interaction between DNA and an Escherichia coli protein omega. J Mol Biol 55, 523–533 (1971).

21. L. F. Liu, C. C. Liu, B. M. Alberts, Type II DNA topoisomerases: enzymes that can unknot a topologically knotted DNA molecule via a reversible double-strand break. Cell 19, 697–707 (1980).

22. A. H. Bizard, I. D. Hickson, The many lives of type IA topoisomerases. J Biol Chem 295, 7138–7153 (2020).

23. Y. Pommier, A. Nussenzweig, S. Takeda, C. Austin, Human topoisomerases and their roles in genome stability and organization. Nat Rev Mol Cell Biol 23, 407–427 (2022).

24. L. K. Saha, Y. Pommier, TOP3A coupling with replication forks and repair of TOP3A cleavage complexes. Cell Cycle 23, 115–130 (2024).

25. C. A. Martin, K. Sarlós, C. V. Logan, R. S. Thakur, D. A. Parry, A. H. Bizard, A. Leitch, L. Cleal, N. S. Ali, M. A. Al-Owain, W. Allen, J. Altmüller, M. Aza-Carmona, B. A. Y. Barakat, J. Barraza-García, A. Begtrup, M. Bogliolo, M. T. Cho, J. Cruz-Rojo, H. A. M. Dhahrabi, N. H. Elcioglu, G. S. Gorman, R. Jobling, I. Kesterton, Y. Kishita, M. Kohda, P. Le Quesne Stabej, A. J. Malallah, P. Nürnberg, A. Ohtake, Y. Okazaki, R. Pujol, M. J. Ramirez, A. Revah-Politi, M. Shimura, P. Stevens, R. W. Taylor, L. Turner, H. Williams, C. Wilson, G. Yigit, L. Zahavich, F. S. Alkuraya, J. Surralles, A. Iglesias, K. Murayama, B. Wollnik, M. Dattani, K. E. Heath, I. D. Hickson, A. P. Jackson, Mutations in TOP3A Cause a Bloom Syndrome-like Disorder. Am J Hum Genet 103, 221–231 (2018).

26. Y. Wang, S. Kaiser, J. Martin-Gonzalez, Y. Li, L. J. Rasmussen, A. J. Lopez-Contreras, A. H. Bizard, I. D. Hickson, Functional analysis of pathological mutations in DNA topoisomerase 3A. Cell Rep 44, 115764 (2025).

27. C. Suski, K. J. Marians, Resolution of converging replication forks by RecQ and topoisomerase III. Mol Cell 30, 779–789 (2008).

28. R. J. DiGate, K. J. Marians, Identification of a potent decatenating enzyme from Escherichia coli. J Biol Chem 263, 13366–13373 (1988).

29. J. Yang, C. Z. Bachrati, J. Ou, I. D. Hickson, G. W. Brown, Human topoisomerase IIIalpha is a single-stranded DNA decatenase that is stimulated by BLM and RMI1. J Biol Chem 285, 21426–21436 (2010).

30. M. Mills, Y. C. Tse-Dinh, K. C. Neuman, Direct observation of topoisomerase IA gate dynamics. Nat Struct Mol Biol 25, 1111–1118 (2018).

31. P. Cejka, J. L. Plank, C. Z. Bachrati, I. D. Hickson, S. C. Kowalczykowski, Rmi1 stimulates decatenation of double Holliday junctions during dissolution by Sgs1-Top3. Nat Struct Mol Biol 17, 1377–1382 (2010).

32. P. Nurse, C. Levine, H. Hassing, K. J. Marians, Topoisomerase III can serve as the cellular decatenase in Escherichia coli. J Biol Chem 278, 8653–8660 (2003).

33. C. M. Lee, G. Wang, A. Pertsinidis, K. J. Marians, Topoisomerase III Acts at the Replication Fork To Remove Precatenanes. J Bacteriol 201, (2019).

34. A. Dadras, A. Le Campion, M. Drolet, Topoisomerase III limits RecA-dependent DNA amplification in the chromosome terminus with RecG. Nucleic Acids Res 54, (2026).

35. J. Brochu, É. Vlachos-Breton, S. Sutherland, M. Martel, M. Drolet, Topoisomerases I and III inhibit R-loop formation to prevent unregulated replication in the chromosomal Ter region of Escherichia coli. PLoS Genet 14, e1007668 (2018).

36. T. D. Deegan, J. Baxter, M. Ortiz Bazán, J. T. P. Yeeles, K. P. M. Labib, Pif1-Family Helicases Support Fork Convergence during DNA Replication Termination in Eukaryotes. Mol Cell 74, 231–244.e239 (2019).

37. K. Mundbjerg, S. W. Jørgensen, J. Fredsøe, I. Nielsen, J. M. Pedersen, I. B. Bentsen, M. Lisby, L. Bjergbaek, A. H. Andersen, Top2 and Sgs1-Top3 Act Redundantly to Ensure rDNA Replication Termination. PLoS Genet 11, e1005697 (2015).

38. L. K. Saha, S. Saha, X. Yang, S. N. Huang, Y. Sun, U. Jo, Y. Pommier, Replication-associated formation and repair of human topoisomerase IIIα cleavage complexes. Nat Commun 14, 1925 (2023).

39. H. Zhang, Y. Sun, S. Saha, L. K. Saha, L. S. Pongor, A. Dhall, Y. Pommier, Genome-wide Mapping of Topoisomerase Binding Sites Suggests Topoisomerase 3α (TOP3A) as a Reader of Transcription-Replication Conflicts (TRC). bioRxiv, 2024.2006.2017.599352 (2024).

40. T. J. Nicholls, C. A. Nadalutti, E. Motori, E. W. Sommerville, G. S. Gorman, S. Basu, E. Hoberg, D. M. Turnbull, P. F. Chinnery, N. G. Larsson, E. Larsson, M. Falkenberg, R. W. Taylor, J. D. Griffith, C. M. Gustafsson, Topoisomerase 3α Is Required for Decatenation and Segregation of Human mtDNA. Mol Cell 69, 9–23.e26 (2018).

41. K. E. Menger, J. Chapman, H. Díaz-Maldonado, M. M. Khazeem, D. Deen, D. Erdinc, J. W. Casement, V. Di Leo, A. Pyle, A. Rodríguez-Luis, I. G. Cowell, M. Falkenberg, C. A. Austin, T. J. Nicholls, Two type I topoisomerases maintain DNA topology in human mitochondria. Nucleic Acids Res 50, 11154–11174 (2022).

42. D. Erdinc, A. Rodríguez-Luis, M. R. Fassad, S. Mackenzie, C. M. Watson, S. Valenzuela, X. Xie, K. E. Menger, K. Sergeant, K. Craig, S. Hopton, G. Falkous, J. Poulton, H. Garcia-Moreno, P. Giunti, C. A. de Moura Aschoff, J. A. Morales Saute, A. J. Kirby, C. Toro, L. Wolfe, D. Novacic, L. Greenbaum, A. Eliyahu, O. Barel, Y. Anikster, R. McFarland, G. S. Gorman, A. M. Schaefer, C. M. Gustafsson, R. W. Taylor, M. Falkenberg, T. J. Nicholls, Pathological variants in TOP3A cause distinct disorders of mitochondrial and nuclear genome stability. EMBO Mol Med 15, e16775 (2023).

43. D. Erdinc, C. A. Albus, A. Rodríguez-Luis, K. E. Menger, A. Thorsell, I. Atanassov, U. Rovšnik, M. Falkenberg, C. M. Gustafsson, T. J. J. Nicholls, Proteolytic cleavage activates the mitochondrial isoform of TOP3A. Nucleic Acids Res 53, (2025).

44. F. B. Couch, C. E. Bansbach, R. Driscoll, J. W. Luzwick, G. G. Glick, R. Bétous, C. M. Carroll, S. Y. Jung, J. Qin, K. A. Cimprich, D. Cortez, ATR phosphorylates SMARCAL1 to prevent replication fork collapse. Genes Dev 27, 1610–1623 (2013).

45. L. I. Toledo, M. Altmeyer, M. B. Rask, C. Lukas, D. H. Larsen, L. K. Povlsen, S. Bekker-Jensen, N. Mailand, J. Bartek, J. Lukas, ATR prohibits replication catastrophe by preventing global exhaustion of RPA. Cell 155, 1088–1103 (2013).

46. J. E. Graham, K. J. Marians, S. C. Kowalczykowski, Independent and Stochastic Action of DNA Polymerases in the Replisome. Cell 169, 1201–1213.e1217 (2017).

47. Y. Baris, M. R. G. Taylor, V. Aria, J. T. P. Yeeles, Fast and efficient DNA replication with purified human proteins. Nature 606, 204–210 (2022).

48. H. Dungrawala, K. L. Rose, K. P. Bhat, K. N. Mohni, G. G. Glick, F. B. Couch, D. Cortez, The Replication Checkpoint Prevents Two Types of Fork Collapse without Regulating Replisome Stability. Mol Cell 59, 998–1010 (2015).

49. B. A. Perez-Cheeks, C. Lee, R. Hayama, K. J. Marians, A role for topoisomerase III in Escherichia coli chromosome segregation. Mol Microbiol 86, 1007–1022 (2012).

50. P. Cejka, J. L. Plank, C. C. Dombrowski, S. C. Kowalczykowski, Decatenation of DNA by the S. cerevisiae Sgs1-Top3-Rmi1 and RPA complex: a mechanism for disentangling chromosomes. Mol Cell 47, 886–896 (2012).

51. M. Chang, M. Bellaoui, C. Zhang, R. Desai, P. Morozov, L. Delgado-Cruzata, R. Rothstein, G. A. Freyer, C. Boone, G. W. Brown, RMI1/NCE4, a suppressor of genome instability, encodes a member of the RecQ helicase/Topo III complex. Embo j 24, 2024–2033 (2005).

52. S. Gangloff, J. P. McDonald, C. Bendixen, L. Arthur, R. Rothstein, The yeast type I topoisomerase Top3 interacts with Sgs1, a DNA helicase homolog: a potential eukaryotic reverse gyrase. Mol Cell Biol 14, 8391–8398 (1994).

53. J. M. Daley, T. Chiba, X. Xue, H. Niu, P. Sung, Multifaceted role of the Topo IIIα-RMI1-RMI2 complex and DNA2 in the BLM-dependent pathway of DNA break end resection. Nucleic Acids Res 42, 11083–11091 (2014).

54. P. Cejka, E. Cannavo, P. Polaczek, T. Masuda-Sasa, S. Pokharel, J. L. Campbell, S. C. Kowalczykowski, DNA end resection by Dna2-Sgs1-RPA and its stimulation by Top3-Rmi1 and Mre11-Rad50-Xrs2. Nature 467, 112–116 (2010).

55. L. Wu, C. Z. Bachrati, J. Ou, C. Xu, J. Yin, M. Chang, W. Wang, L. Li, G. W. Brown, I. D. Hickson, BLAP75/RMI1 promotes the BLM-dependent dissolution of homologous recombination intermediates. Proc Natl Acad Sci U S A 103, 4068–4073 (2006).

56. D. Xu, R. Guo, A. Sobeck, C. Z. Bachrati, J. Yang, T. Enomoto, G. W. Brown, M. E. Hoatlin, I. D. Hickson, W. Wang, RMI, a new OB-fold complex essential for Bloom syndrome protein to maintain genome stability. Genes Dev 22, 2843–2855 (2008).

57. L. Wu, I. D. Hickson, The Bloom’s syndrome helicase suppresses crossing over during homologous recombination. Nature 426, 870–874 (2003).

58. A. H. Bizard, J. F. Allemand, T. Hassenkam, M. Paramasivam, K. Sarlós, M. I. Singh, I. D. Hickson, PICH and TOP3A cooperate to induce positive DNA supercoiling. Nat Struct Mol Biol 26, 267–274 (2019).

59. J. L. Plank, S. H. Chu, J. R. Pohlhaus, T. Wilson-Sali, T. S. Hsieh, Drosophila melanogaster topoisomerase IIIalpha preferentially relaxes a positively or negatively supercoiled bubble substrate and is essential during development. J Biol Chem 280, 3564–3573 (2005).

60. J. Walter, L. Sun, J. Newport, Regulated chromosomal DNA replication in the absence of a nucleus. Mol Cell 1, 519–529 (1998).

61. E. J. Vontalge, T. Kavlashvili, S. N. Dahmen, M. T. Cranford, J. M. Dewar, Control of DNA replication in vitro using a reversible replication barrier. Nat Protoc 19, 1940–1983 (2024).

62. R. A. Laskey, A. D. Mills, N. R. Morris, Assembly of SV40 chromatin in a cell-free system from Xenopus eggs. Cell 10, 237–243 (1977).

63. D. T. Gruszka, S. Xie, H. Kimura, H. Yardimci, Single-molecule imaging reveals control of parental histone recycling by free histones during DNA replication. Sci Adv 6, (2020).

64. J. Walter, J. Newport, Initiation of eukaryotic DNA replication: origin unwinding and sequential chromatin association of Cdc45, RPA, and DNA polymerase alpha. Mol Cell 5, 617–627 (2000).

65. R. Hanai, P. R. Caron, J. C. Wang, Human TOP3: a single-copy gene encoding DNA topoisomerase III. Proc Natl Acad Sci U S A 93, 3653–3657 (1996).

66. W. Li, S. M. Kim, J. Lee, W. G. Dunphy, Absence of BLM leads to accumulation of chromosomal DNA breaks during both unperturbed and disrupted S phases. J Cell Biol 165, 801–812 (2004).

67. L. V. Campos, S. X. Van Ravenstein, E. J. Vontalge, B. H. Greer, D. R. Heintzman, T. Kavlashvili, W. H. McDonald, K. L. Rose, B. F. Eichman, J. M. Dewar, RTEL1 and MCM10 overcome topological stress during vertebrate replication termination. Cell Rep 42, 112109 (2023).

68. T. Han, M. Goralski, E. Capota, S. B. Padrick, J. Kim, Y. Xie, D. Nijhawan, The antitumor toxin CD437 is a direct inhibitor of DNA polymerase α. Nat Chem Biol 12, 511–515 (2016).

69. A. Ercilla, J. Benada, S. Amitash, G. Zonderland, G. Baldi, K. Somyajit, F. Ochs, V. Costanzo, J. Lukas, L. Toledo, Physiological Tolerance to ssDNA Enables Strand Uncoupling during DNA Replication. Cell Rep 30, 2416–2429.e2417 (2020).

70. K. P. M. Mehta, V. Thada, R. Zhao, A. Krishnamoorthy, M. Leser, K. Lindsey Rose, D. Cortez, CHK1 phosphorylates PRIMPOL to promote replication stress tolerance. Sci Adv 8, eabm0314 (2022).

71. V. Aria, J. T. P. Yeeles, Mechanism of Bidirectional Leading-Strand Synthesis Establishment at Eukaryotic DNA Replication Origins. Mol Cell 73, 199–211.e110 (2018).

72. J. K. Barrows, D. T. Long, Cell-free transcription in Xenopus egg extract. J Biol Chem 294, 19645–19654 (2019).

73. T. E. T. Mevissen, M. Kümmecke, E. W. Schmid, L. Farnung, J. C. Walter, STK19 positions TFIIH for cell-free transcription-coupled DNA repair. Cell 187, 7091–7106.e7024 (2024).

74. K. L. Chan, T. Palmai-Pallag, S. Ying, I. D. Hickson, Replication stress induces sister-chromatid bridging at fragile site loci in mitosis. Nat Cell Biol 11, 753–760 (2009).

75. T. R. Singh, A. M. Ali, V. Busygina, S. Raynard, Q. Fan, C. H. Du, P. R. Andreassen, P. Sung, A. R. Meetei, BLAP18/RMI2, a novel OB-fold-containing protein, is an essential component of the Bloom helicase-double Holliday junction dissolvasome. Genes Dev 22, 2856–2868 (2008).

76. J. Yin, A. Sobeck, C. Xu, A. R. Meetei, M. Hoatlin, L. Li, W. Wang, BLAP75, an essential component of Bloom’s syndrome protein complexes that maintain genome integrity. Embo j 24, 1465–1476 (2005).

77. N. A. Ellis, J. Groden, T. Z. Ye, J. Straughen, D. J. Lennon, S. Ciocci, M. Proytcheva, J. German, The Bloom’s syndrome gene product is homologous to RecQ helicases. Cell 83, 655–666 (1995).

78. R. Bermejo, T. Capra, V. Gonzalez-Huici, D. Fachinetti, A. Cocito, G. Natoli, Y. Katou, H. Mori, K. Kurokawa, K. Shirahige, M. Foiani, Genome-organizing factors Top2 and Hmo1 prevent chromosome fragility at sites of S phase transcription. Cell 138, 870–884 (2009).

79. E. Yi, R. Chamorro González, A. G. Henssen, R. G. W. Verhaak, Extrachromosomal DNA amplifications in cancer. Nat Rev Genet 23, 760–771 (2022).

80. R. Lebofsky, T. Takahashi, J. C. Walter, DNA replication in nucleus-free Xenopus egg extracts. Methods Mol Biol 521, 229–252 (2009).

81. J. M. Dewar, M. Budzowska, J. C. Walter, The mechanism of DNA replication termination in vertebrates. Nature 525, 345–350 (2015).

82. H. Goulaouic, T. Roulon, O. Flamand, L. Grondard, F. Lavelle, J. F. Riou, Purification and characterization of human DNA topoisomerase IIIalpha. Nucleic Acids Res 27, 2443–2450 (1999).

