## Supplementary figures and images for "Topoisomerase IIIα resolves inter- and intra-molecular intertwines during DNA replication"

### Supplementary Figures

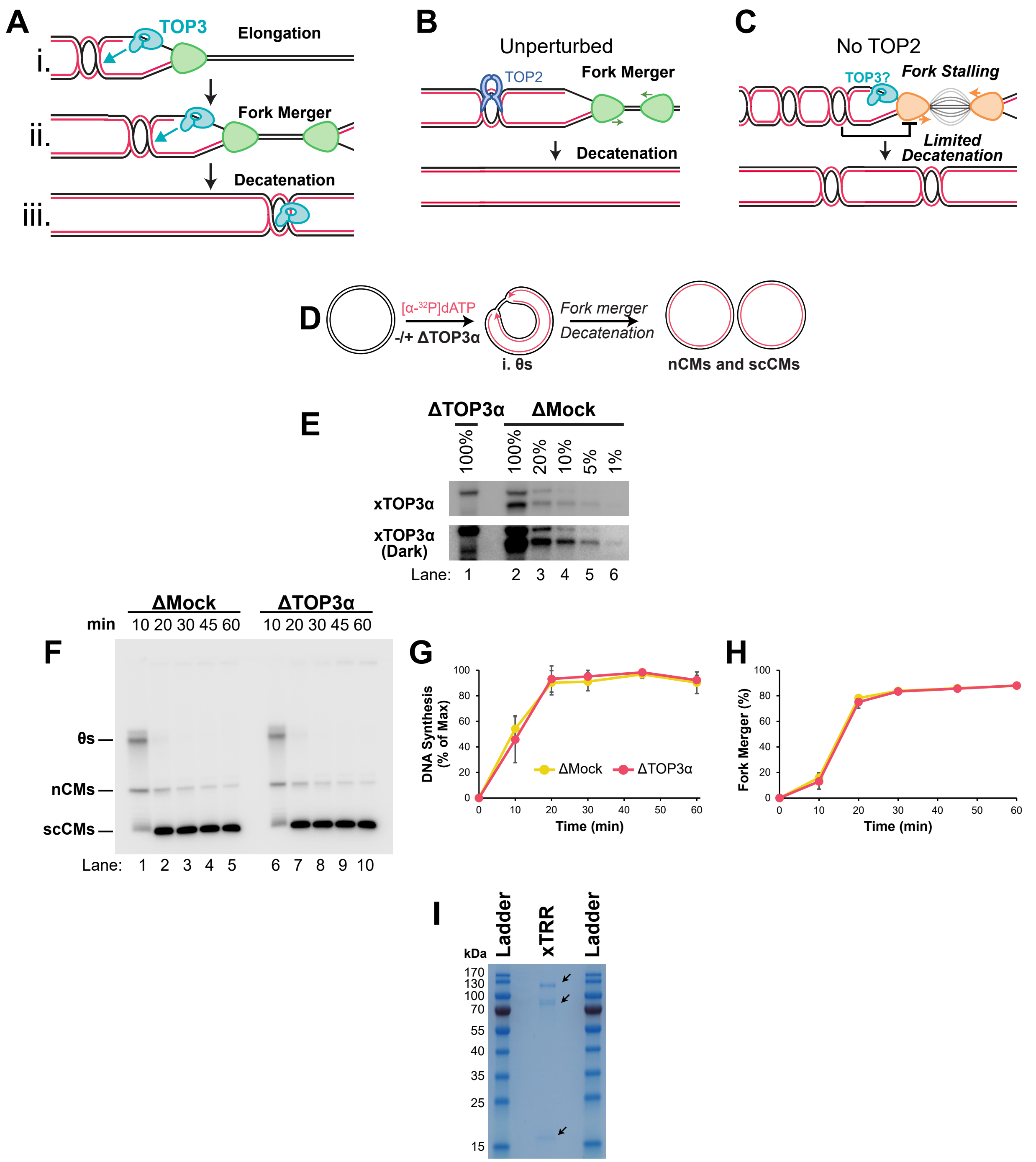

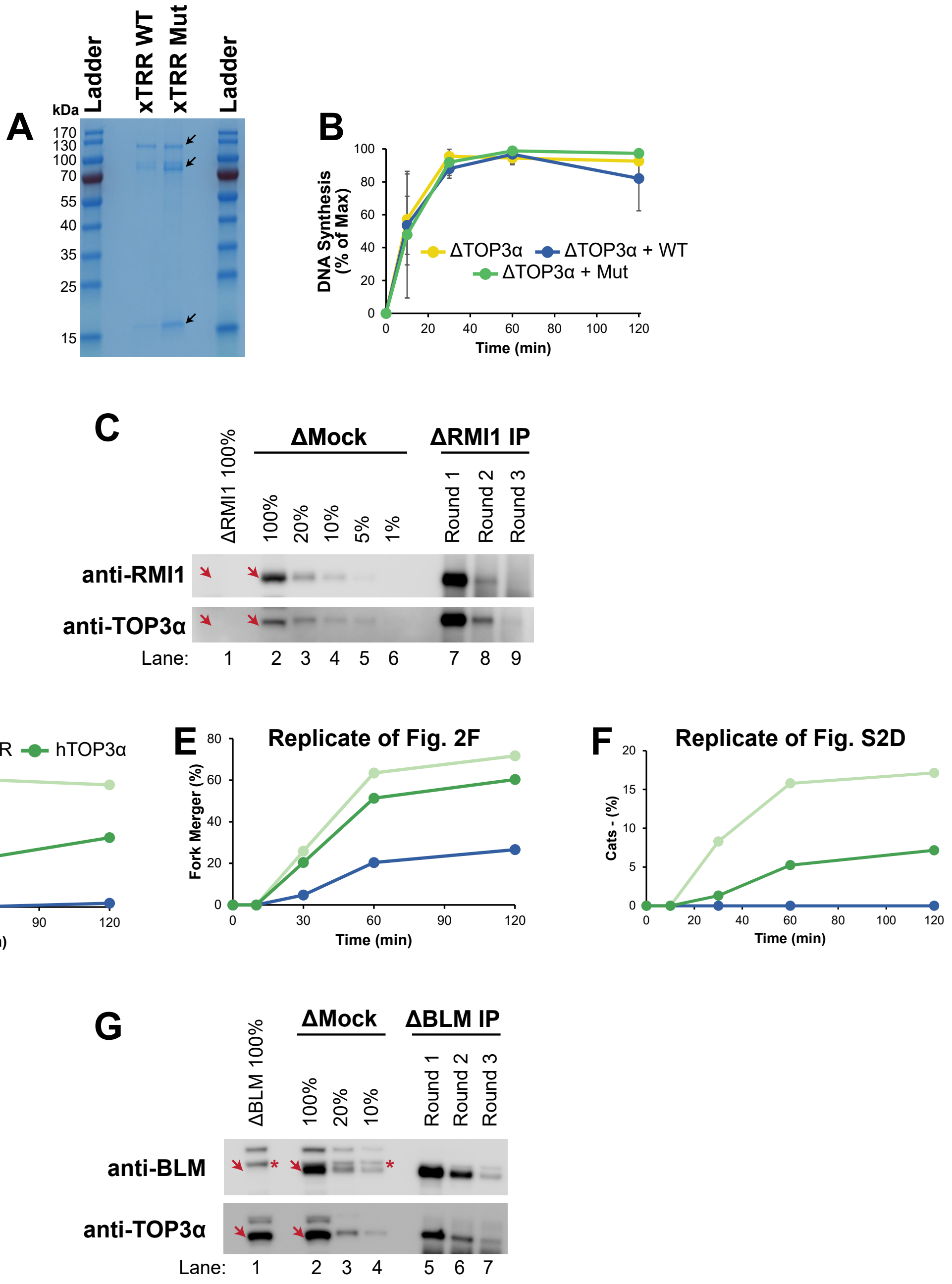

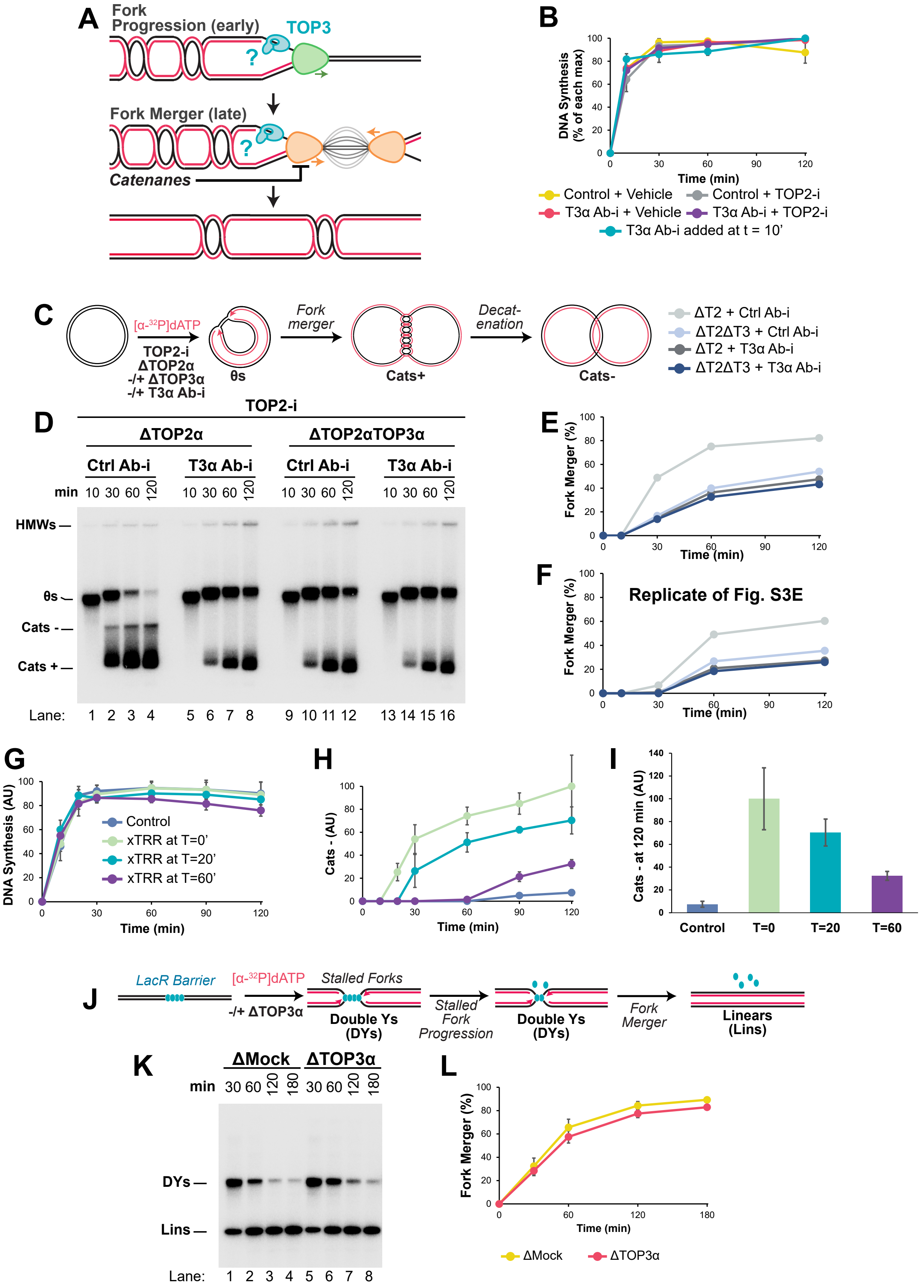

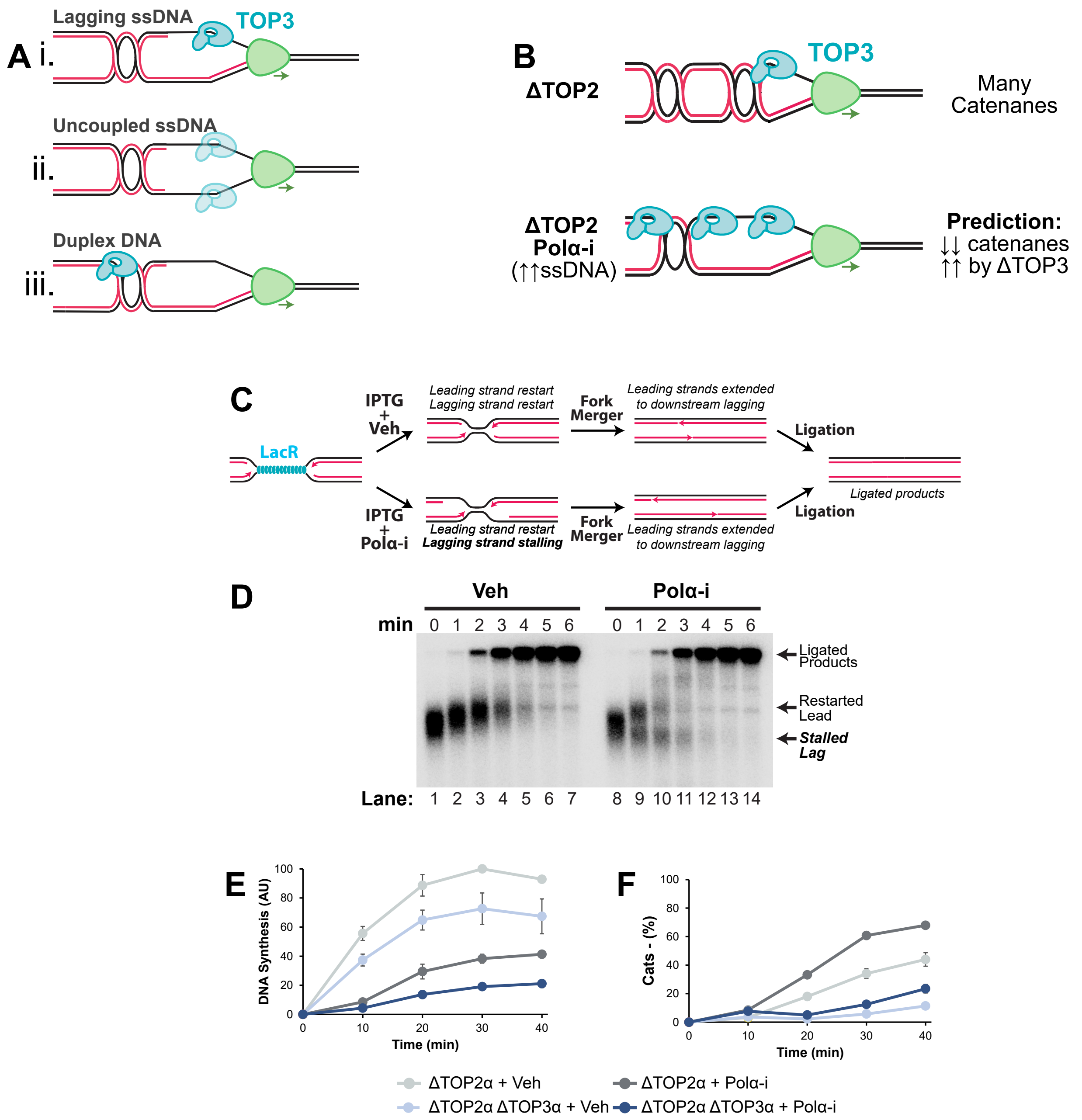

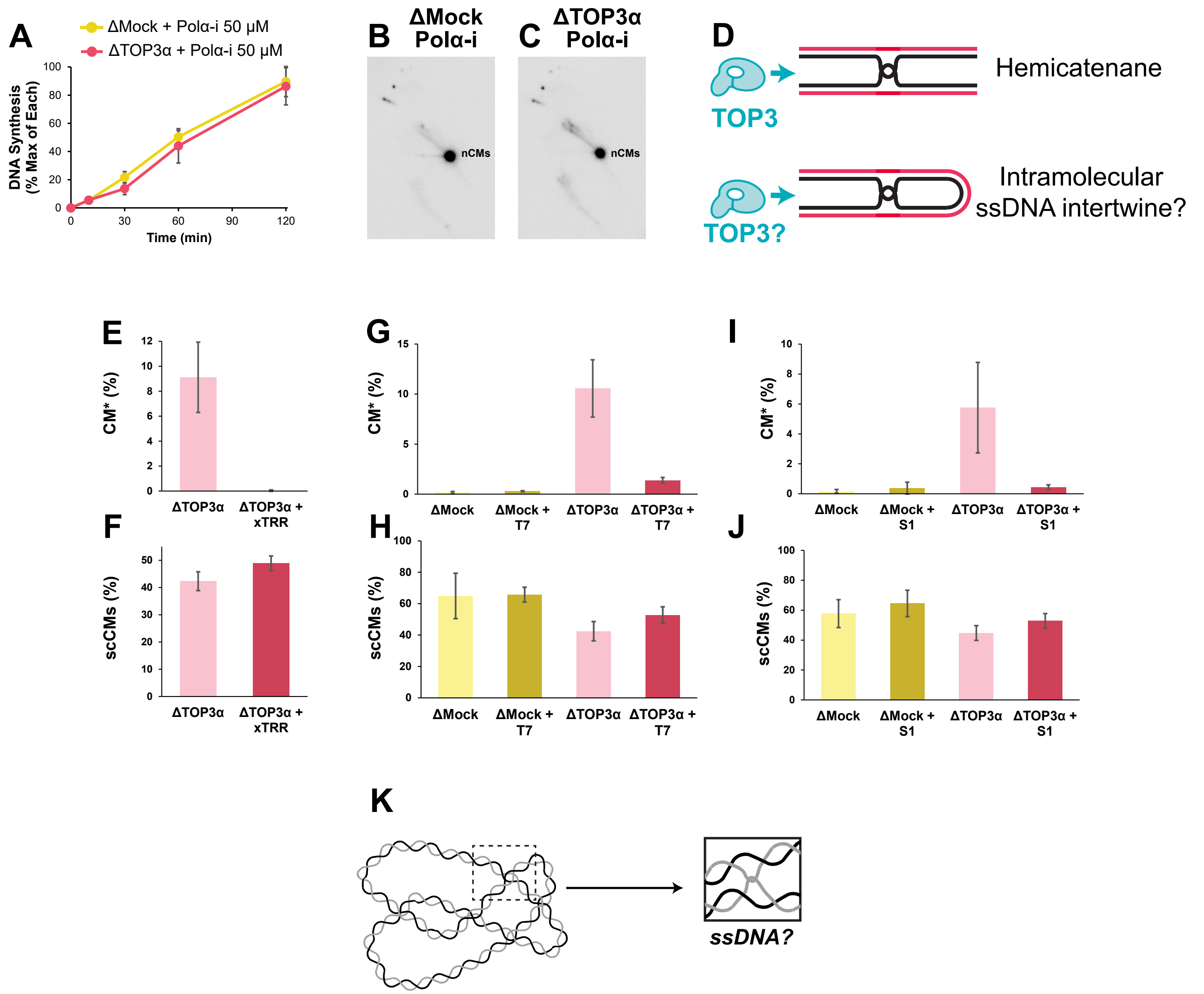
